# H3-K27M-Mutant Nucleosomes Interact with MLL1 to Shape the Glioma Epigenetic Landscape

**DOI:** 10.1101/2021.08.09.455680

**Authors:** Noa Furth, Danielle Algranati, Bareket Dassa, Olga Beresh, Vadim Fedyuk, Natasha Morris, Lawryn H. Kasper, Dan Jones, Michelle Monje, Suzanne J. Baker, Efrat Shema

**Affiliations:** Department of Biological Regulation, Weizmann Institute of Science, Rehovot 76100, Israel; Bioinformatics Unit, Department of Life Sciences Core Facilities, Faculty of Biochemistry, Weizmann Institute of Science; Rehovot 76100, Israel; Department of Developmental Neurobiology, St. Jude Children’s Research Hospital, Memphis, TN 38105, USA; SeqLL, Woburn, MA 01801, USA; Department of Neurology, Stanford University School of Medicine, Stanford, CA 94305, USA

## Abstract

Cancer-associated mutations in genes encoding histones dramatically reshape chromatin and support tumorigenesis. Lysine to methionine substitution of residue 27 on histone H3 (K27M) is a driver mutation in high-grade pediatric gliomas, known to abrogate Polycomb Repressive Complex 2 (PRC2) activity. We applied single-molecule systems to image individual nucleosomes and delineate the combinatorial epigenetic patterns associated with H3-K27M expression. We found that chromatin marks on H3-K27M-mutant nucleosomes are dictated both by their incorporation preferences and by intrinsic properties of the mutation. Mutant nucleosomes not only preferentially bind PRC2, but also directly interact with MLL1, thus leading to genome-wide redistribution of H3K4me3. H3-K27M-mediated deregulation of both repressive and active chromatin marks leads to unbalanced ‘bivalent’ chromatin, which may support a poorly differentiated cellular state. This study provides evidence for a direct effect of H3-K27M oncohistone on the MLL1-H3K4me3 pathway and highlights the capability of single-molecule tools to reveal mechanisms of chromatin deregulation in cancer.

## Introduction

Chromatin structure regulates cell-type specific transcriptional programs to establish cellular identity (Rivera and Ren, 2013). Nucleosomes, the fundamental building block of chromatin, are extensively modified by covalent attachment of various chemical groups (Cavalli and Heard, 2019; Schuettengruber et al., 2017). While an individual epigenetic mark may have specific roles, many studies support the notion of a complex epigenetic network that requires combinatorial presence of several marks to control biological functions (Berger, 2007). This network of modifications plays vital roles in cellular differentiation, and is disrupted in many types of cancers, facilitating tumor initiation, progression and resistance to therapy (Flavahan et al., 2017; Valencia and Kadoch, 2019; Zhao and Shilatifard, 2019).

One such example is the frequent deregulation of PRC2, which catalyzes the addition of di- and tri-methylation of lysine 27 on histone H3 (H3K27me2/3) (Schuettengruber et al., 2017). Mutations in histone proteins, such as substitution of H3K27 with methionine (H3-K27M mutation), also abrogate PRC2 activity and are frequent in childhood high-grade gliomas. In diffuse intrinsic pontine glioma (DIPG), an aggressive and fatal malignancy, up to 80% of diagnosed children exhibit H3-K27M mutations, which can occur in genes encoding either the H3.3 variant or canonical H3.1/3.2 (Castel et al., 2015; Fontebasso et al., 2014; Schwartzentruber et al., 2012; Wu et al., 2012). These mutant histones, estimated to represent only a minor fraction of the chromatin H3 pool, induce global and dramatic changes to the epigenetic landscape. Seminal works have shown global loss of H3K27me3 in H3-K27M cells, with the exact mechanism mediating this loss still under debate (Bender et al., 2013a; Chan et al., 2013; Lewis et al., 2013b). Biochemical and structural studies revealed tight binding of the mutant histone to EZH2, perhaps leading to its sequestration (Justin et al., 2016; Lewis et al., 2013b). Nevertheless, Chromatin Immunoprecipitation and sequencing (ChIP-seq) studies do not support aberrant localization of Polycomb complex to H3-K27M enriched regions (Mohammad et al., 2017; Piunti et al., 2017). Recently, Brien and colleagues showed that H3.3-K27M incorporation is associated with increased PRC1/2 binding in pre-existing Polycomb target sites (Brien et al., 2021). Furthermore, there is evidence that EZH2 is persistently inhibited by H3-K27M even upon its dissociation from mutant nucleosomes (Lee et al., 2019; Stafford et al., 2018), and H3-K27M mutant-like peptides can abrogate PRC2 activity not within a chromatin context (Jain et al., 2019; Piunti et al., 2019). The extent of this inhibition, and its potential effects on PRC2 enzymatic and spreading activities, is under active research (Harutyunyan et al., 2020; Harutyunyan et al., 2019; Jain et al., 2020b). Along with global loss of H3K27me3, H3-K27M is associated with a prominent increase in open chromatin marks such as H3K27ac and H3K36me3 (Krug et al., 2019; Lewis et al., 2013a; Piunti et al., 2017). Overall, H3-K27M histones were suggested to exert both *cis* effects on modifications within the mutant nucleosome, and *trans* effects on wild-type (WT) nucleosomes (Lewis et al., 2013b; Weinberg et al., 2017; Yu et al., 2019). However, the mechanisms by which low prevalence H3-K27M nucleosomes exert oncogenic pressure remain limited and controversial.

In this study, we applied single-molecule epigenetic profiling technology to systematically decode combinatorial modification patterns of individual WT and H3-K27M-mutant nucleosomes in pediatric gliomas. We provide evidence for direct deregulation of two critical epigenetic pathways, Polycomb and Trithorax, by mutant nucleosomes, with global and local ramifications to the epigenetic landscape and expression programs of glioma cells.

## Results

### Single-molecule imaging and quantification of epigenetic alterations induced by H3.1-K27M and H3.3-K27M mutant nucleosomes

We recently described a single-nucleosome imaging platform that allows direct counting and decoding of modified nucleosomes (Shema et al., 2016), thus providing a unique tool to study how H3-K27M mutant histones affect the epigenetic landscape. Cell-derived mono-nucleosomes are anchored in a spatially distributed manner on PEG-coated surface. Captured nucleosomes are incubated with fluorescently-labelled antibodies directed against specific histone modifications. Total Internal Reflection (TIRF) microscopy is utilized to record the position and modification state of each nucleosome. Time series images are taken, to allow detection of maximal binding events, and to account for varying binding kinetics of different antibodies (Fig. 1A, S1A). As demonstrated with recombinant nucleosomes, the single-molecule system is highly sensitive and quantitative, allowing detection of minute fractions of mutant nucleosomes within a mixed population (Fig. S1B). Furthermore, accumulating data from multiple images taken over time results in detection of maximal antibody binding events, as seen from saturation of the signal (Fig. S1C).

**Figure 1:**
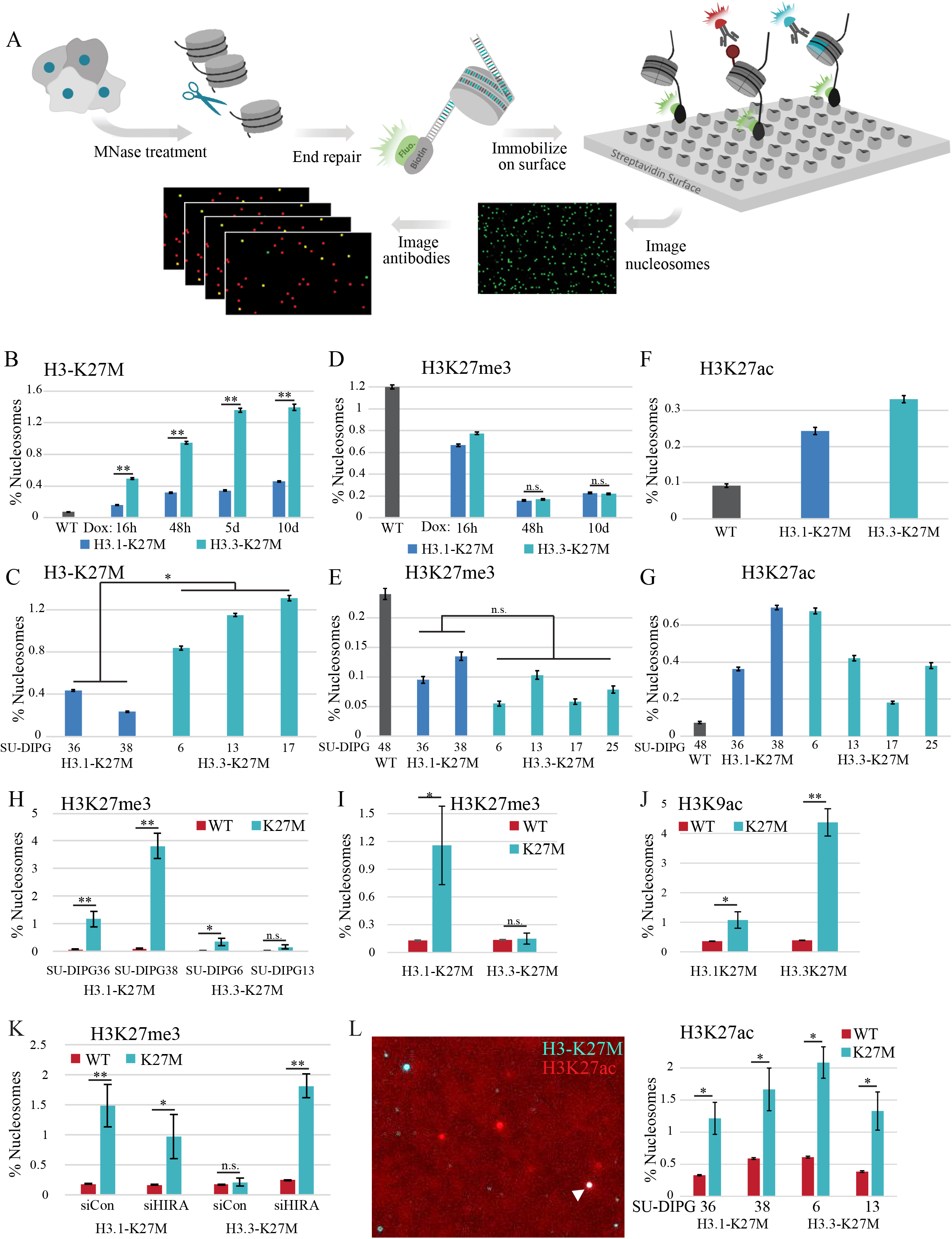
Single-molecule imaging of H3-K27M nucleosomes reveals induction of robust epigenetic programs. **(A)** Scheme of the single-molecule imaging experimental setup. **(B-C)** Single-molecule imaging and quantification of H3-K27M-mutant nucleosomes in HEK293 cells induced to express H3.1-K27M or H3.3-K27M **(B)** and in different patient-derived DIPG cultures **(C)**. H3.3-K27M shows higher chromatin incorporation than H3.1-K27M. *p-val<0.05, **p-val<0.001. Dox = Doxycycline. **(D-G)** Single-molecule imaging of H3K27me3 **(D-E)** and H3K27ac **(F-G)** in the inducible HEK293 system **(D, F)** and the patient derived DIPG cultures **(E, G)**, similar to B-C. Both H3.1-K27M and H3.3-K27M variants induce robust reduction in H3K27me3 and an increase in H3K27ac. **(H-I)** Single-molecule analysis of H3K27me3 on WT or H3-K27M-mutant nucleosomes in the indicated DIPG lines **(H)** and in HEK293 cells expressing H3.1-K27M or H3.3-K27M **(I).** Strong enrichment of H3K27me3 is seen specifically on H3.1-K27M-mutant nucleosomes. **(J)** Single-molecule analysis of H3K9ac on WT or H3-K27M-mutant nucleosomes in HEK293 cells expressing mutant H3. Strong enrichment of H3K9ac is seen specifically on H3.3-K27M mutant nucleosomes. **(K)** HEK293 cells were transfected with the indicated siRNAs and the expression of either H3.1-K27M or H3.3-K27M histones was induced by doxycycline. Nucleosomes from the indicated samples were analyzed as in H. Depletion of HIRA alters H3.3 incorporation and results in enrichment of H3K27me3 also on H3.3-K27M-mutant nucleosomes. **(L)** Single-molecule analysis of H3K27ac on H3-K27M-mutant nucleosomes in the indicated DIPG lines. Left panel: a representative image of individual nucleosomes imaged with TIRF. White arrow points to a H3-K27M-H3K27ac asymmetric nucleosome. Right panel: percentage of wild type (WT) or H3-K27M-mutant nucleosomes acetylated on lysine 27. Enrichment of H3K27ac on mutant nucleosomes (i.e. heterotypic nucleosomes) is observed for both variants.

We first took advantage of this system to explore chromatin incorporation dynamics of H3-K27M-mutant nucleosomes. To generate a controlled expression system, we infected HEK293 cells with an inducible construct encoding either H3.1- or H3.3-K27M gene. Interestingly, despite being expressed from similar constructs, H3.3-K27M protein levels are higher than H3.1-K27M, and show increased levels of chromatin incorporation (Fig. 1B, S2A-C). Similarly, analysis of a panel of patient-derived DIPG cultures expressing the H3-K27M mutation endogenously, revealed higher levels of mutant nucleosomes in cells carrying K27M mutation on H3.3 compared with canonical H3.1-K27M (Fig. 1C, S2D).

Despite the observed differences in protein and incorporation levels of the H3.1- and H3.3-K27M-mutant nucleosomes, both led to robust and global decrease in H3K27me3, as well documented by others (Bender et al., 2013a; Lee et al., 2019; Lewis et al., 2013b). In fact, the fraction of H3K27me3 nucleosomes measured by single-molecule is highly similar between H3.3-K27M and H3.1-K27M cells, in both the inducible system and the DIPG lines, and is significantly lower than H3K27me3 in WT cells (Fig. 1D-E). Along with the loss of the repressive H3K27me3 mark, we observed an increase in histone acetylation (H3K27ac and H3K9ac, Fig. 1F-G, S3), in agreement with previous reports (Lewis et al., 2013b; Piunti et al., 2017). Overall, this data validates the single-molecule methodology for quantitative analysis of epigenetic alterations, and suggests a threshold effect for H3-K27M-nucleosomes in mediating global loss of H3K27me3 and gain of H3K27ac, which is not directly correlated with H3-K27M levels.

We next used Cut&Run (Skene and Henikoff, 2017) in the patient-derived DIPG cultures to study the genomic distribution of mutant nucleosomes. We found that in cells expressing H3.3-K27M (SU-DIPG6, SU-DIPG13 and SU-DIPG17), the genomic locations of H3-K27M are similar between the lines, and mimic the distribution of WT H3.3 (Fig. S4). On the contrary, the signal of mutant nucleosomes from H3.1-K27M cells (SU-DIPG36 and SU-DIPG38) does not aggregate to distinct peaks, although comparable number of reads and fragment size were obtained (Fig S4B-C and Table S1). This is likely due to the unbiased replication-coupled deposition of canonical H3.1 across the genome (Tagami et al., 2004), and the limitation of assays that rely on averaging signals over a population of cells. These results are in agreement with previous reports showing that H3-K27M mutation does not affect genome-wide distribution of the H3 variants (Brien et al., 2021; Nagaraja et al., 2019; Sarthy et al., 2020).

### Decoding asymmetric Lysine 27 modifications on H3-K27M nucleosomes

Our single-molecule imaging platform has the inherent advantage of decoding combinations of marks on individual nucleosomes by multiplexing specific antibodies, thus resolving epigenetic features of H3-K27M-mutant versus WT nucleosomes. H3-K27M mutation has been shown to alter EZH2 chromatin binding and kinetics (Fig. S5A) (Bender et al., 2013a; Jain et al., 2020a; Leicher et al., 2020; Lewis et al., 2013b; Tatavosian et al., 2018). Indeed, we observed that upon expression of H3-K27M, EZH2 genomic distribution is altered towards distal genomic regions (Fig S5B). We aimed to further explore this interaction by probing whether individual mutant nucleosomes are marked by H3K27me3. To that end, we scored all nucleosomes as either K27M or WT, and quantified the percentage of nucleosomes that harbor H3K27me3 in each group. While canonical H3.1-mutant nucleosomes are highly enriched for H3K27me3 (generating asymmetric H3.1-K27M-K27me3 structures), only mild enrichment is observed for H3.3-mutant nucleosomes (Fig. 1H-I). On the contrary, the active acetylation mark on histone H3 lysine 9 (H3K9ac) showed marked preference for H3.3-mutant nucleosomes, likely reflecting its association with active regions (Fig. 1J and S5C). These results support the notion that H3-K27M variants, similar to WT H3 variants, are incorporated to different genomic regions and thus acquire different chromatin modifications.

The enrichment of H3K27me3 on H3.1-K27M nucleosomes may result from preferential binding of PRC2 and local methylation of the WT H3 within mutant nucleosomes. We further speculated that H3.3-mutant nucleosomes, which are preferentially incorporated to euchromatic regions by the HIRA chaperone complex (Goldberg et al., 2010), might accumulate active modifications that block PRC2 activity (Stafford et al., 2018; Yu et al., 2019), thus explaining the reduced enrichment of H3K27me3. In order to test this hypothesis, we knocked down the H3.3-specific histone chaperone HIRA that is responsible for H3.3 deposition in promoters, gene bodies and active regulatory elements (Fig. S5D) (Goldberg et al., 2010). Of note, H3.3 is also incorporated by the ATRX-DAXX complex to closed chromatin regions such as telomeres and pericentromeric heterochromatin. Thus, depletion of HIRA is expected to shift the balance of H3.3 deposition from active to closed regions (Goldberg et al., 2010; Wong et al., 2010). As expected, silencing of HIRA led to overall reduction in chromatin incorporation of H3.3-K27M, but not H3.1-K27M, validating the role of this chaperone in depositing WT and mutant H3.3 (Fig. S5E). Moreover, we observed a decrease in H3.3-K27M-H3K9ac nucleosomes, supporting a reduction of H3.3 nucleosomes incorporated to open chromatin regions (Fig. S5F). Interestingly, while in control cells H3K27me3 is only enriched on H3.1-mutant nucleosomes, upon HIRA depletion we observed a marked increase in H3.3-K27M-H3K27me3 nucleosomes, supporting the notion that these nucleosomes are preferentially formed in closed chromatin regions (Fig. 1K).

Our Cut&Run data (Fig. S4B) and previous reports^17^ suggest that mutant nucleosomes can be asymmetrically acetylated on lysine 27 of the WT histone H3, forming heterotypic nucleosome complexes. Indeed, H3-K27M nucleosomes are acetylated at higher prevalence than WT nucleosomes, providing unequivocal evidence for the presence of heterotypic nucleosome complexes (Fig. 1L and S5G-H). To test whether this enrichment is only due to incorporation preference of H3.3 to open regions, we compared acetylation specifically on the WT H3.3 variant versus the mutant H3.3 by exogenously expressing FLAG-tagged versions of these histones and probing with a combination of H3K27ac and anti-FLAG antibodies. Albeit mild, the H3.3-K27M nucleosomes show stronger enrichment for H3K27ac as compared to WT H3.3 nucleosomes (Fig. S5I). Furthermore, the association between H3-K27M and H3K27ac was also observed for the canonical H3.1-K27M, raising the intriguing possibility that it reflects not only the incorporation preference of the H3.3 variant to open regions, but also a mechanistic link between mutant nucleosomes and lysine 27 acetylation. Of note, pervasive deposition of H3K27ac mark in K27M-mutant cells has been recently suggested (Brien et al., 2021; Krug et al., 2019), which may be driven, at least partially, by this preferred association. Overall, our single-molecule analysis suggests that chromatin modifications present on K27M-mutant nucleosomes are dictated by both the chromatin environment to which each variant is incorporated, and intrinsic properties of the mutant histone, which may alter the binding and activity of chromatin modifiers.

### H3-K27M-mutant nucleosomes are enriched for H3K4me3 and preferentially bind MLL1

To identify additional *cis* and *trans* effects of H3-K27M, we applied our single-molecule method to systematically quantify various epigenetic marks on WT and H3-K27M nucleosomes. Interestingly, combinatorial probing for H3-K27M and H3K4me3 revealed marked enrichment of H3K4me3 on both H3.1- and H3.3-mutant nucleosomes (Fig. 2A-B, Fig S6A). While the H3.3-K27M variant might be expected to co-localize with H3K4me3 due to its deposition to promoters, the significant enrichment of H3K4me3 also on the canonical mutant histone (H3.1-K27M) suggests a mechanism independent of incorporation preferences. Of note, H3-K27M expression did not profoundly alter global levels of H3K4me3-modified nucleosomes (Fig. 2C, S6B). To further explore the association between mutant nucleosomes and H3K4me3, we performed Cut&Run in patient-derived DIPG cultures, containing the H3.3-K27M mutation. Indeed, H3-K27M peaks co-localized with H3K4me3 signal; importantly, this association was not restricted to peaks around promoter regions (accounting for only ~20% of H3.3-K27M peaks, Fig. S7A) and was seen also for H3-K27M peaks annotated in distal genomic regions (Fig. 2D-E, S7B-C). As H3.1-K27M Cut&Run reads did not aggregate to peaks (Fig. S4C), we could not perform similar analysis for H3.1. The single-molecule data showing H3K4me3 enrichment for both H3.1-K27M and H3.3-mutant nucleosomes in the HEK293 inducible isogenic system and the patient-derived glioma cells, as well as the enrichment of H3K4me3 on H3.3-K27M nucleosomes at distal regions, support a potential mechanistic link between mutant nucleosomes and H3K4me3.

**Figure 2:**
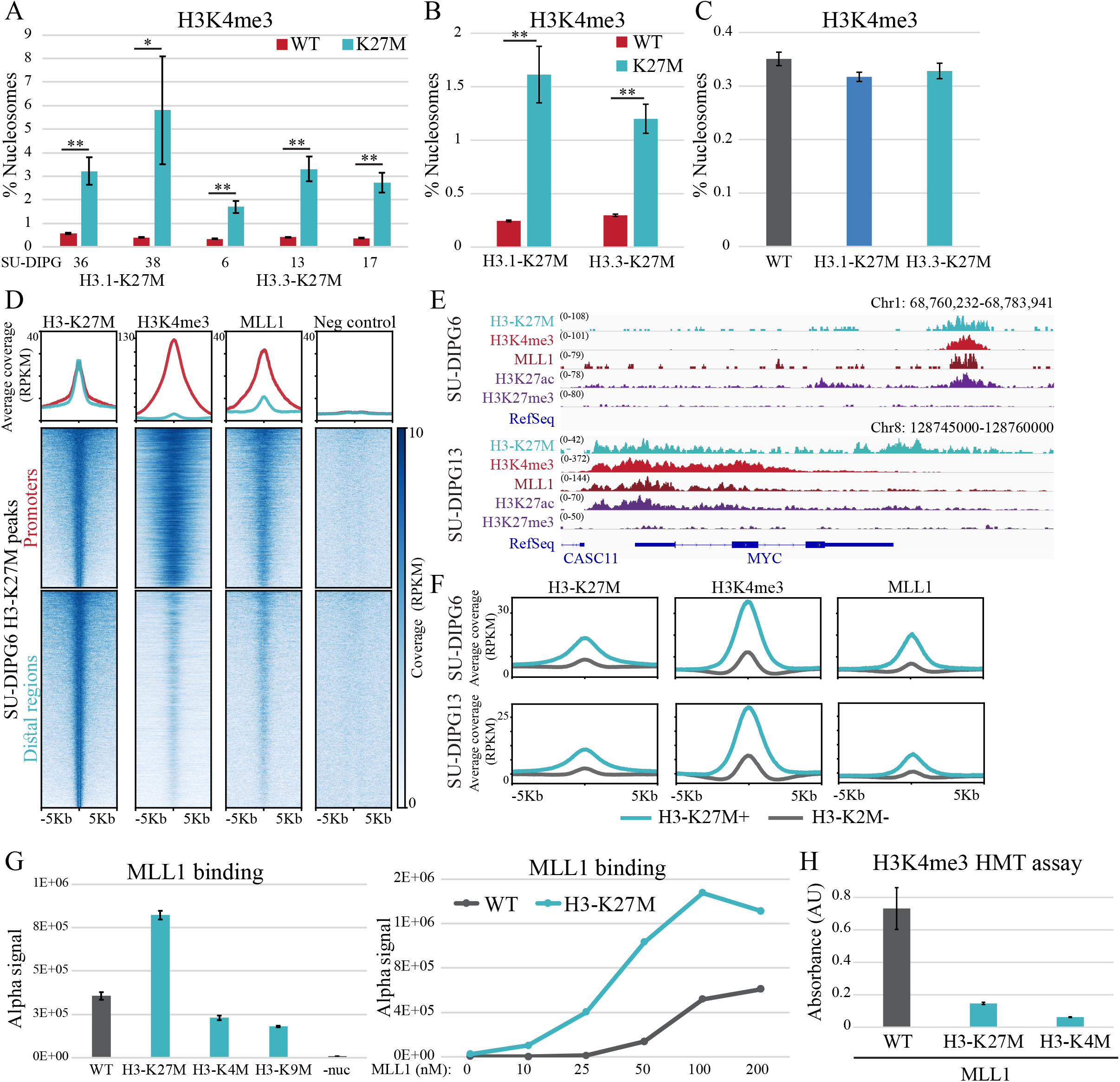
H3-K27M-mutant nucleosomes directly recruit MLL1 and are enriched for H3K4me3. **(A-B)** Single-molecule analysis of H3K4me3 on WT or H3-K27M-mutant nucleosomes in the indicated DIPG cultures **(A)** and in HEK293 cells expressing H3.1-K27M or H3.3-K27M **(B)**. Both H3.1-K27M and H3.3-K27M nucleosomes show high and similar enrichment for H3K4me3. *p-val<0.05, **p-val<0.001. **(C)** Single-molecule quantification of H3K4me3 nucleosomes in the HEK293 cells expressing either WT H3, H3.1-K27M or H3.3-K27M. **(D)** H3-K27M peaks identified in SU-DIPG6 cells are associated with H3K4me3 and MLL1 in both promoters and distal genomic regions. Heatmap shows H3-K27M peaks annotated to promoters (±3Kb from TSS, 11,274 peaks) or distal genomic regions (15,337 peaks) using ChIPseeker algorithm sorted in descending order from top to bottom (center represents the peak summit, left and right borders represent −5kb and +5kb, respectively). Signal for H3K4me3, MLL1 and negative control antibody (anti-rabbit 2^nd^ antibody) in these genomic regions is plotted accordingly. Average coverage for each group is plotted on top. **(E)** Representative IGV genome browser tracks of regions enriched for H3-K27M, H3K4me3 and MLL1. Shown are intergenic region (top) and the MYC gene (bottom). **(F)** H3K27ac peaks which do not overlap with TSS (±2Kb) were defined as enhancers and divided according to the presence of H3-K27M peaks. Average coverages of MLL1, H3K4me3 and H3-K27M are plotted for each group in either SU-DIPG6 (upper panel) or SU-DIPG13 (bottom panel). H3-K27M peaks are enriched for MLL1 binding also at enhancer regions. **(G)** AlphaLISA protein-protein binding assay indicates preferential binding of MLL1 to H3-K27M-mutant nucleosomes over WT nucleosomes. Left panel: 50nM of GST-tagged MLL1 was incubated with 10nM of biotinylated recombinant nucleosomes. Alpha acceptor and donor beads were added, and the Alpha signal corresponding to protein-protein binding was measured. Average ± SD of two technical replicates are shown. Right panel: varying amounts of GST-tagged MLL1 were incubated with 10nM biotinylated recombinant nucleosomes and protein-protein interaction were measured as in the upper panel. See methods and Fig. S8C for details about the AlphaLISA assay. **(H)** The indicated recombinant nucleosomes were incubated with 0.2μg recombinant KMT2A (MLL1) complex and H3K4me3 methylation was measured. Mean ± SD of two technical repeats is shown. Higher global methylation level is detected for WT nucleosomes versus H3-K27M nucleosome.

H3K4 trimethylation is mediated by methyltransferases of the mixed-lineage leukemia (MLL) family members, mainly MLL1 and MLL2, as well as SET Domain Containing 1A and 1B (SET1A and SET1B) (Schuettengruber et al., 2017). MLL1 is known to deposit H3K4me3 on developmental genes, is recurrently deregulated in different tumors (Rao and Dou, 2015), and robustly expressed in DIPG (Filbin et al., 2018). Similar to H3K4me3, Cut&Run analysis of DIPG cells indicated that H3-K27M peaks were enriched for MLL1 binding, in both promoter and distal regions (Fig. 2D-E, S7A). In fact, about 60% of the identified MLL1 peaks co-localized with H3-K27M peaks (Fig. S8A), and this association is stronger on H3-K27M positive peaks compared to H3.3 peaks which do not show H3-K27M signal (Fig. S8B). Enhancers containing K27M-mutant nucleosomes showed higher H3K4me3 and MLL1 signals compared to enhancers without H3-K27M, providing further evidence for preferential interaction between MLL1 and H3-K27M nucleosomes, not limited to transcription start site (TSS) proximal regions (Fig. 2F).

These findings led us to hypothesize a potential direct interaction between MLL1 and H3-K27M, perhaps mimicking the increased binding affinity between EZH2 and H3-K27M nucleosomes (Bender et al., 2013a; Lewis et al., 2013b). As both enzymes contain a SET domain, MLL1 may favor binding to the mutant histone over the WT, as observed for EZH2 (Fig. S5A) (Bender et al., 2013a; Lewis et al., 2013b). To test our hypothesis, we performed a protein-protein interaction assay between recombinant GST-tagged MLL1 complex and WT or H3-K27M biotinylated recombinant nucleosomes (AlphaLISA assay, S8C and methods). Strikingly, we observed increased binding between H3-K27M-mutant nucleosomes and MLL1 complex, indicating direct preferential interaction (Fig. 2G, Fig. S8D). Moreover, *in-vitro* methylation assay revealed that recombinant MLL1 is less effective in methylation of lysine 4 when incubated with H3-K27M mutant nucleosomes compared to WT nucleosomes (Fig. 2H). Taken together, these results strongly support increased binding of MLL1 to mutant nucleosomes. This preferential binding may promote local K4 trimethylation of H3-K27M nucleosomes, but also reduce its effective methylation of a large amount of target molecules. Although not strongly affecting global levels of H3K4me3 in cells (Fig. 2C), possibly due to additional compensating methyl-transferases, this aberrant interaction may alter the regulation of this active chromatin mark.

### H3K4me3 is aberrantly distributed in H3-K27M mutant cells

To explore whether aberrant binding of MLL1 to K27M-mutant nucleosomes may affect the deposition patterns of H3K4me3, we examined the genome-wide patterns of this mark in the DIPG lines. To that end, we analyzed H3K4me3 peaks which are present in both WT and mutant H3 cells (shared, >80% overlap), and unique H3K4me3 peaks which are only present in H3-K27M cells (Fig. 3A, S9A). As expected, both H3.3 and H3-K27M are associated with H3K4me3 signal, due to the incorporation of this variant at promoters. We focused on H3K4me3 peaks that are specific to the K27M-mutant cells (unique), and are highly enriched for H3-K27M (Fig. 3A). Interestingly, we found these peaks to be depleted from promoter regions in comparison to H3K4me3 unique peaks devoid of H3-K27M (Fig. 3A-B, S9B). This may suggest an aberrant distribution of H3K4me3 to non-promoter regions, mediated by MLL1 association with H3-K27M. Indeed, total H3K4me3 peaks in H3.3-K27M-mutant cells show reduced enrichment with gene promoters; the main genomic feature associated with H3K4me3, compared to WT H3 cells (Fig. 3C).

**Figure 3:**
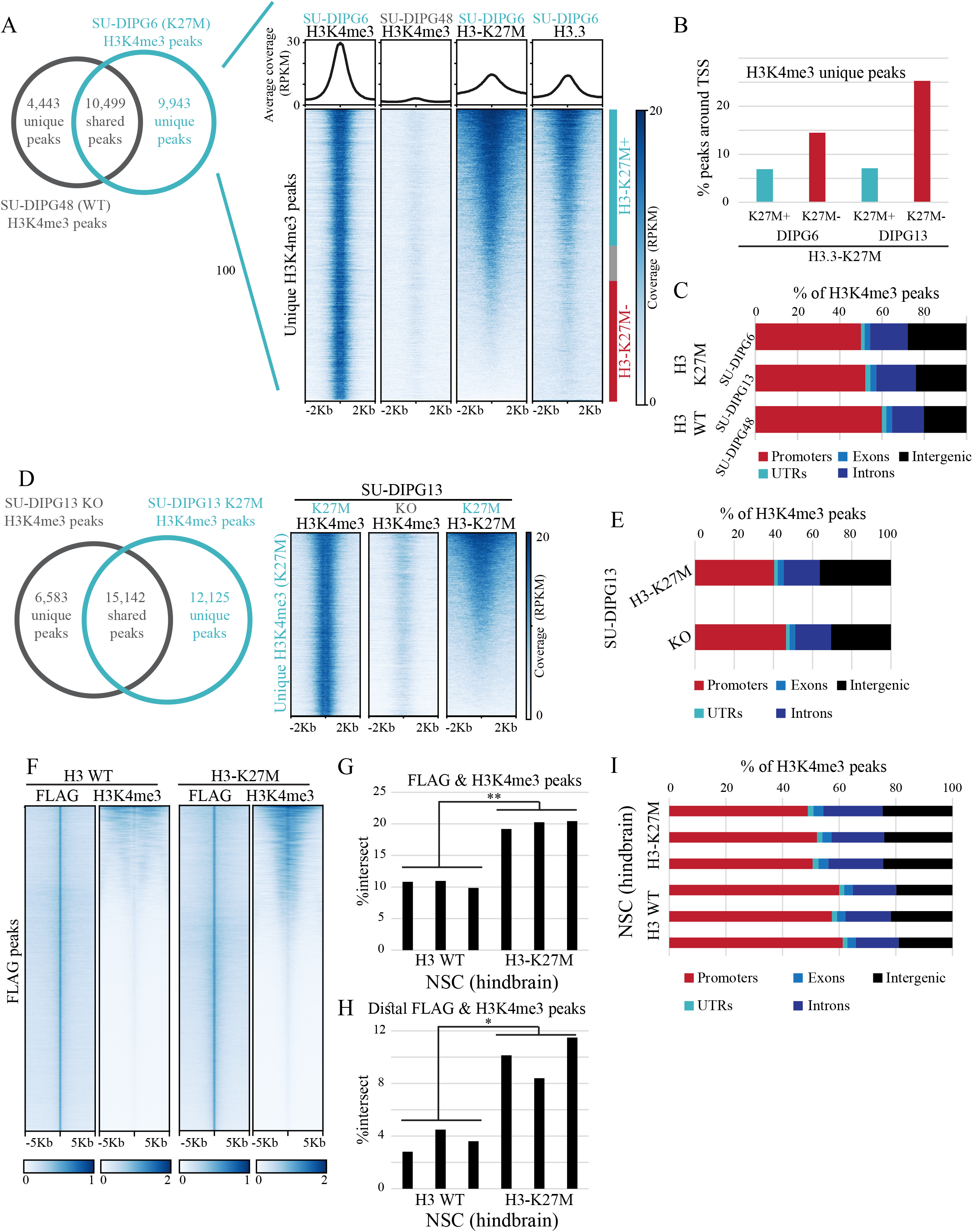
Aberrant distribution of H3K4me3 in H3.3-K27M mutant cells. **(A)** Unique H3K4me3 peaks in SU-DIPG6 are highly enriched for H3-K27M. Left: Venn diagram of H3K4me3 peaks in WT (SU-DIPG48) compared to H3-K27M-mutant (SU-DIPG6) cell lines. Unique peaks are defined as peaks that do not overlap between the lines; shared peaks are defined as peaks with minimum of 80% overlap. Right: Heatmap represents H3K4me3 peaks that are unique to SU-DIPG6 (9,943 peaks) with the corresponding H3K4me3 signal in WT SU-DIPG48 (center represents the peak summit, left and right borders represent −5kb and +5kb, respectively). H3-K27M and H3.3 signal in these genomic regions is plotted accordingly. Regions are annotated according to the overlap with H3-K27M peak (bar on the right); regions with at least 80% overlap (blue) or no overlap (red). Heatmap is sorted according to annotation. Average coverage is plotted on top. **(B)** H3K4me3 unique peaks (see panel B) were divided according to their overlap with H3-K27M peaks and the percentage of peaks within ±5Kb form the TSS is shown. H3K4me3 peaks that are associated with H3-K27M are depleted from TSS regions. **(C)** Fraction of H3K4me3 peaks that correspond to the indicated genomic features in the indicated cultures. WT H3 cells have higher fraction of H3K4me3 peaks associated with promoters. **(D)** Left: venn diagram of H3K4me3 peaks in K27M-mutant compared to K27M-KO isogenic cell lines. Unique peaks are defined as peaks that do not overlap between the lines; shared peaks are defined as peaks with minimum of 80%overlap. Right: Heatmap represents H3K4me3 peaks that are unique to H3.3-K27M cells. H3-K27M signal in these genomic regions is plotted accordingly. H3K4me3 peaks unique to K27M cells are enriched for H3-K27M signal. **(E)** H3K4me3 peaks in K27M-KO or control DIPG cultures were analyzed as in D. A smaller fraction of H3K4me3 is associated with promoters in cells expressing H3-K27M. **(F)** ChIP-seq using FLAG and H3K4me3 antibodies in hindbrain NSC expressing FLAG-tagged WT or K27M-mutant H3.3 reveal high association of H3K4me3 specifically with K27M-mutant nucleosomes. Heatmap of FLAG peaks detected in one WT and one mutant sample shown along with the corresponding H3K4me3 signal. Heatmap shows ±5kb around the peak summit, and sorted according to H3K4me3 signal. Average coverage is plotted on top. **(G)** Percentage of FLAG peaks that overlap with H3K4me3 peaks (minimal overlap of 25%) from three mice from each genotype. **p-val<0.001 **(H)** Higher fraction of H3K4me3 peaks are associated with FLAG signal at distal genomic elements in H3-K27M cells. Percentage of H3K4me3 peaks that overlap with FLAG peaks, annotated to distal genomic regions, is shown (n=3 for each genotype). *p-val<0.01 **(I)** H3K4me3 peaks identified in WT or H3.3-K27M hindbrain NSC were analyzed as in D.

To validate these results, we repeated the analysis above in an isogenic SU-DIPG13 cells in which the H3.3-K27M gene has been knocked-out (KO) (Harutyunyan et al., 2019; Krug et al., 2019). Analysis of H3K4me3 peaks unique to the cells expressing H3.3-K27M confirmed their association with the mutant histone (Fig. 3D, S9C). In fact, H3.3-K27M cells show a larger number of unique H3K4me3 peaks, and more than 50% of these peaks are associated with H3-K27M. Furthermore, knock-out of the mutant histone resulted in higher MLL1 signal on transcription start sites (TSS, Fig. S9D) and increased the fraction of H3K4me3 peaks which is associated with gene promoters (Fig. 3E). Thus, the recruitment of MLL1 by K27M-mutant nucleosomes may facilitate aberrant deposition of H3K4me3 and an altered epigenetic landscape.

To further confirm the functional association between H3-K27M and H3K4me3 in an additional isogenic model, we took advantage of the powerful mouse system generated by Larson and colleagues (Larson et al., 2019). This system allows direct comparison between the WT and K27M-mutant histone, due to the conditional expression of an engineered *H3f3a* gene resulting in expression of FLAG-HA tagged H3.3 or H3.3-K27M. Strikingly, ChIP-seq for FLAG and H3K4me3 in hindbrain neural stem cells (NSC) from these mice revealed strong association of H3K4me3 specifically with the mutant histone (Fig. 3F-G, S9E). Furthermore, the percentage of H3K4me3 peaks that overlap with FLAG peaks found in distal genomic regions is higher in the cells expressing the mutant histone (Fig. 3H). Finally, as seen in human DIPG cultures, expression of the mutant histone reduces the fraction of H3K4me3 that is associated with gene promoters (Fig. 3I).

### Deregulation of H3K4me3 is associated with mutant-specific transcriptional programs

We further explored the functional outcomes of H3-K27M-MLL1 interaction by profiling chromatin accessibility in the DIPG cultures by Assay for Transposase Accessible Chromatin followed by sequencing (ATAC-seq) (Buenrostro et al., 2015). Focusing on H3K4me3-unique peaks (described in fig. 3A), we observed higher accessibility in H3K4me3 regions associated with H3-K27M (Fig. 4A). As expected, the H3-K27M positive peaks also showed increased binding of MLL1, supporting a functional role for the H3-K27M-MLL1 interaction. To assess whether this interaction affects gene expression, we profiled the transcriptome of several DIPG cultures by RNA-seq. In line with the more accessible chromatin, genes that are associated with H3-K27M peaks (out of the H3K4me3-unique regions) are expressed at higher levels (Fig. 4B). Of note, this is despite the smaller fraction of these H3K4me3-K27M+ peaks that localize to promoter regions (Fig. 3B). Thus, the MLL1-H3-K27M interaction results in aberrant deposition of H3K4me3 and higher chromatin accessibility in both genic and non-genic regions. When occurring on genes, it is associated with increased expression.

**Figure 4:**
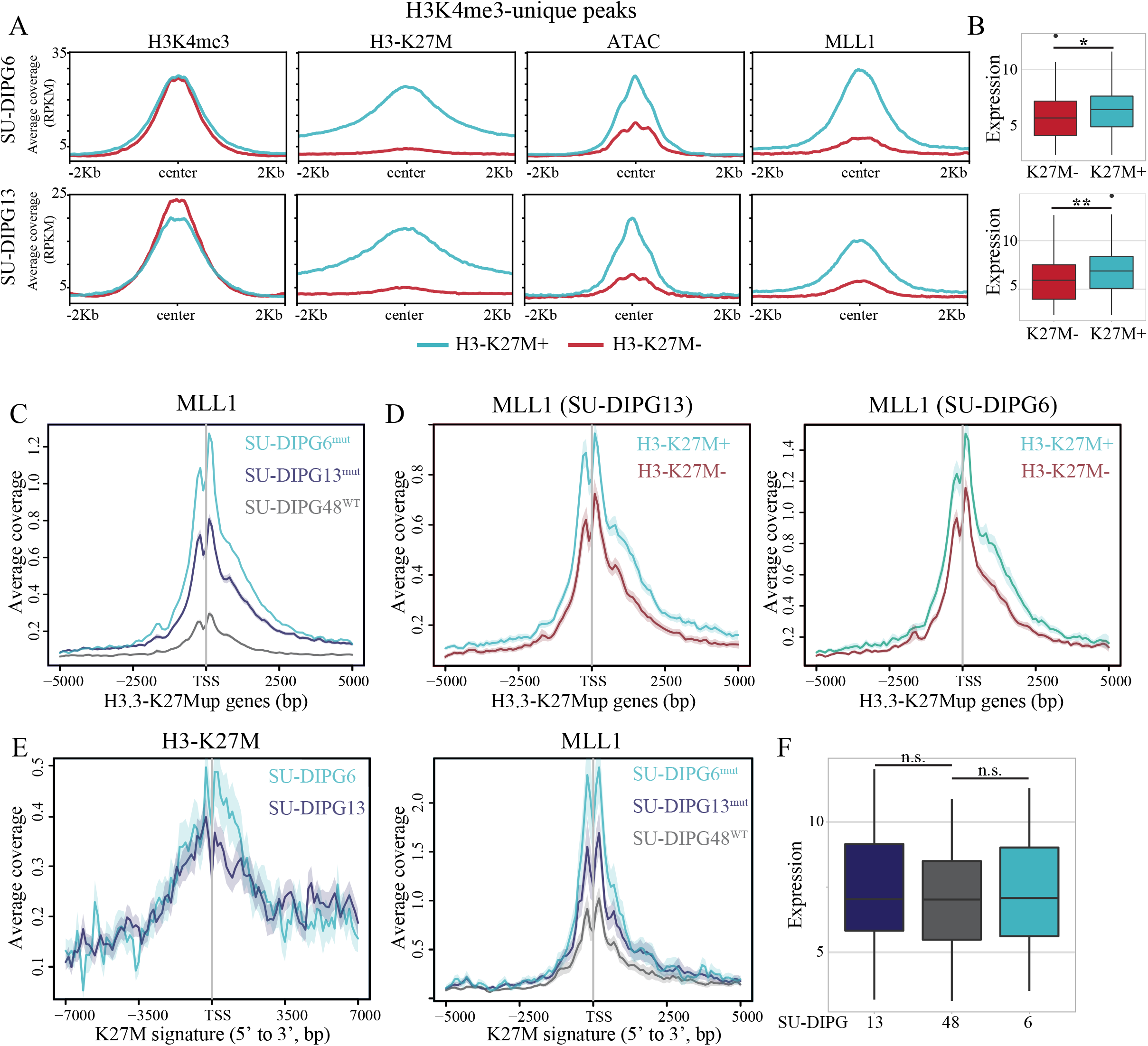
H3-K27M interaction with MLL1 contributes to mutant-specific transcriptional programs. **(A)** H3K4me3 unique peaks (as defined in fig. 4B) were divided according to their overlap with H3-K27M peaks. Average coverage of H3K4me3, H3-K27M, ATAC and MLL1 in each group is shown for SU-DIPG6 (upper panel) and SU-DIPG13 (bottom panel). H3K4me3 peaks that are associated with H3-K27M are more accessible and enriched for MLL1 binding. **(B)** Average expression levels (log2) of genes associated with the groups of peaks shown in A. Higher expression levels are seen for genes that associate with the unique H3K4me3-H3-K27M+ peaks. *p-val<0.01, **p-val<0.001. **(C)** MLL1 coverage (read count per million mapped reads) ±5Kb from the TSS of genes upregulated in H3.3-K27M cells (n=1079 genes, see also Fig. S10A-B). H3.3-K27M upregulated genes are enriched for MLL1 binding compared to H3 WT cells. **(D)** H3.3-K27M upregulated genes were divided according to the presence of H3-K27M peak ±2Kb form their TSS. MLL1 coverage in SU-DIPG13 (left) and SU-DIPG6 (right) is shown for each group. Higher MLL1 coverage is seen for genes harboring H3-K27M peak around their TSS. **(E)** H3-K27M (left) and MLL1 (right) coverage around TSSs of genes comprising H3-K27M-specific signature from single-cell transcriptomic analysis of gliomas, from Filbin et al.(Filbin et al., 2018) (n=80 genes). H3-K27M-mutant signature is associated with higher coverage of MLL1 in H3-K27M cells (SU-DIPG6 and SU-DIPG13) compared to H3 WT cells (SU-DIPG48). **(F)** Average expression levels (log2) of the H3-K27M signature shown in **(E)** in the indicated DIPG cultures.

Next, we analyzed the expression data to define a transcriptional signature upregulated in cells expressing H3.3-K27M (H3.3-K27Mup, Fig. S10A-B). This signature is strongly associated with MLL1 signal in both SU-DIPG6 and SU-DIPG13 (H3.3-K27M), compared to SU-DIPG48 (H3 WT, Fig. 4C). Further division of this signature according to the signal of H3-K27M reveals higher MLL coverage associated with genes that contain H3-K27M at their promoters, supporting the increased binding between the mutant histone and MLL1 (Fig. 4D). Of note, the H3-K27M+ genes are expressed at higher levels (Fig. S10C), which may either stem from the increased interaction with MLL1, or due to preferential incorporation of H3.3 nucleosomes to transcribed regions (Ahmad and Henikoff, 2002). To address this point, we took advantage of a published signature of genes from a single-cell RNA-seq study suggested to associate with H3-K27M in tumor samples (Filbin et al., 2018). Indeed, we found H3-K27M deposited in the TSS region of these genes in both SU-DIPG6 and SU-DIPG13. Importantly, these genes also showed an increase in MLL1 binding in H3.3-mutant cells compared to WT, despite similar averaged expression levels in our cultures (Fig. 4E-F). Overall, our results suggest that the H3-K27M-MLL1 interaction promotes aberrant distribution of H3K4me3, which may contribute to mutant-specific transcriptional programs.

### Residual H3K27me3 nucleosomes are enriched for bivalency in H3-K27M cells

The repressive mark H3K27me3 and the active mark H3K4me3, both aberrantly distributed in cells expressing H3-K27M-nucleosomes, can co-exist to generate bivalent nucleosomes (Bernstein et al., 2006). This bivalent state, initially identified in stem cells to mark promoters of key developmental genes, is also associated with cancer and indicates a poorly differentiated state (Béguelin et al., 2013; Shema et al., 2016; Zhao and Shilatifard, 2019). Bivalency is suggested to play important roles in tumor plasticity and drug resistance (Brown et al., 2014). In DIPG, recent work demonstrated aberrant activation of bivalent genes associated with tumor development (Larson et al., 2019).

When analyzing the genome-wide profiles of H3K4me3 and H3K27me3 in patient-derived H3-K27M DIPG cells, we observed that, while there is a global loss of H3K27me3 peaks, the number of bivalent regions remains similar (Fig. 5A-C). This enrichment of bivalent regions was seen also in the human isogenic SU-DIPG13 cells (Fig. 5D). We applied single-molecule imaging to probe for bivalent (H3K27me3+ H3K4me3+) nucleosomes extracted from HEK293 cells expressing WT or mutant H3, as well as from SU-DIPG13 cells expressing endogenous H3.3-K27M or knocked-out for this allele (KO). Despite drastic global reduction in H3K27me3 nucleosomes in H3-K27M expressing cells, the residual H3K27me3 nucleosomes were highly enriched for H3K4me3, resulting in bivalency (Fig. 5E-F, Fig. S11A-B).

**Figure 5:**
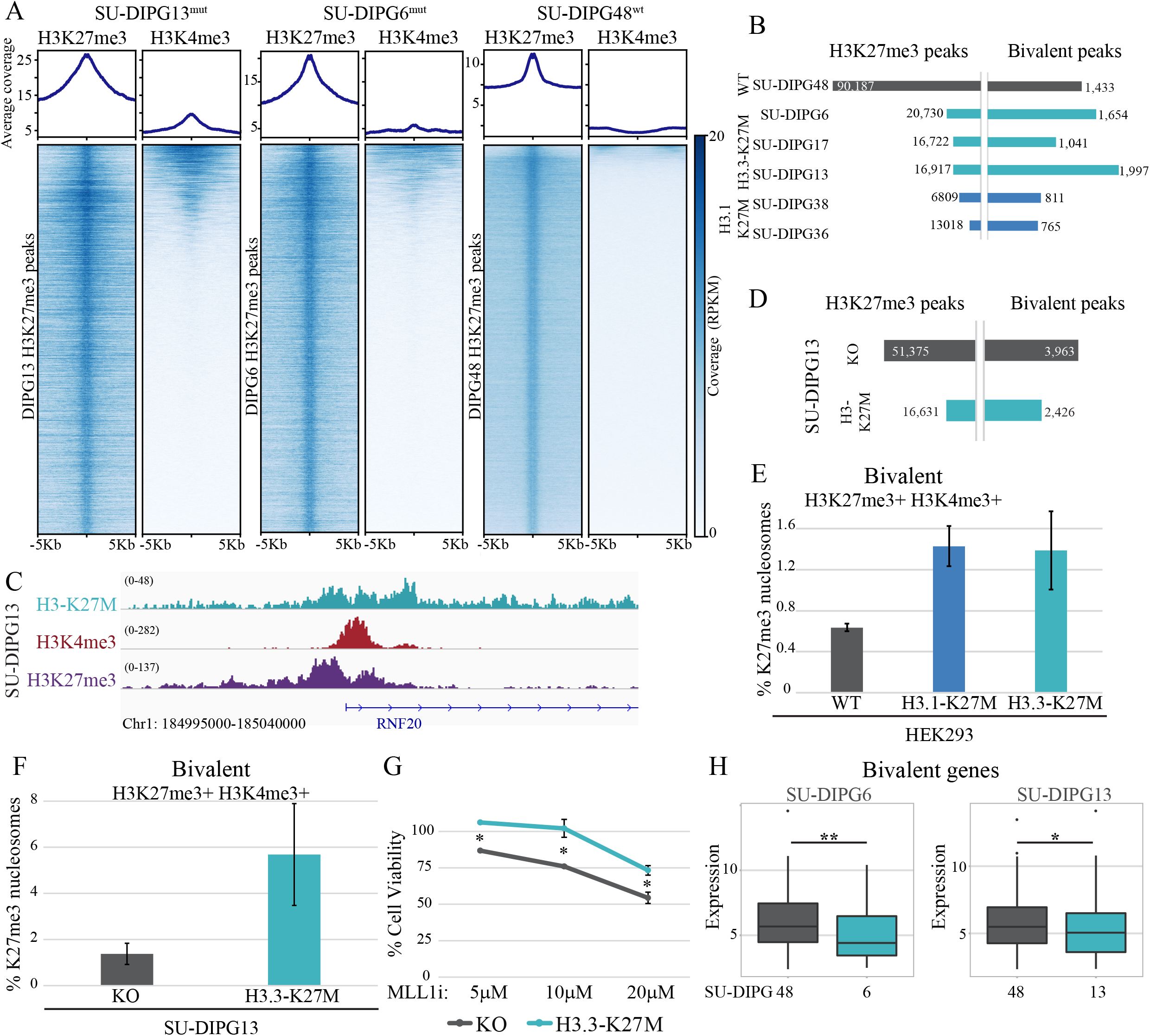
Residual H3K27me3 nucleosomes show higher levels of bivalency. **(A)** Heatmap of H3K27me3 peaks in SU-DIPG13 (H3-K27M, 16,722 peaks), SU-DIPG6 (H3K27M, 20,730) and SU-DIPG48 (H3-WT, 90,187), with the corresponding H3K4me3 signal in descending order. High fraction of bivalent regions is observed in H3-K27M cells. **(B)** Number of H3K27me3 peaks and bivalent peaks identified by Cut&Run analysis in the indicated samples. Bivalent peaks are defined as H3K4me3 peaks with at least 80% overlap with H3K27me3. **(C)** Reads’ coverage on representative bivalent loci in H3-K27M-mutant lines. **(D)** Number of H3K27me3 peaks and bivalent peaks identified by Cut&Run analysis of H3-K27M and KO DIPG culture (SU-DIPG13). Bivalent peaks were defined as in B. **(E)** Single-molecule analysis of bivalent nucleosomes in HEK293 cells expressing either WT H3, H3.1-K27M or H3.3-K27M. Shown is the percentage of bivalent nucleosomes out of H3K27me3 nucleosomes. Average ± SE of at least two experiments is shown. Expression of H3-K27M leads to higher percentage of bivalent nucleosomes, despite the global loss of H3K27me3 nucleosomes. **(F)** Single-molecule analysis of bivalent nucleosomes in the isogenic DIPG culture (SU-DIPG13) expressing the endogenous H3-K27M or knocked-out for the K27M-mutant allele (KO). **(G)** SU-DIPG13 knocked out for H3-K27M (KO) and their corresponding control cells were treated with MLL1 inhibitor (MM-102) at the indicated concentrations. Cell viability was measured by CellTiterGlo assay relative to DMSO treated cells (mean ± SE for three technical replicates, *p-val<0.05). Cells lacking H3-K27M are more sensitive to MLL1 inhibition. **(H)** Average expression levels (log2) of genes associated with the bivalent regions (±3Kb from TSS) identified in SU-DIPG6 (n=763, left) and SU-DIPG13 (n=1003, right) are shown for each of the DIPG lines. Bivalent genes are expressed at lower levels in K27M-mutant cells compared to WT. *p-val<0.05, **p-val<0.001

To further explore the functional significance of these two epigenetic pathways to DIPG biology, we treated the SU-DIPG13 isogenic system with MLL1 and EZH2 inhibitors. In agreement with previous reports, pharmacological inhibition of EZH2 in SU-DIPG13 cells resulted in significant cell death, indicating that residual H3K27me3 in H3-K27M cells is vital for cells’ survival (Mohammad et al., 2017). Cells knocked-out for K27M showed reduced sensitivity to EZH2 inhibition, in line with their higher H3K27me3 levels (Fig. S11C). Interestingly, an opposite effect is seen upon MLL1 inhibition; cells lacking H3-K27M (SU-DIPG13 KO cells) showed higher sensitivity to this inhibitor, albeit the effects are mild (Fig. 5G). Similar trend is seen when comparing SU-DIPG48 to SU-DIPG13: cells expressing H3-K27M are less sensitive to MLL1 inhibition (Figure S11D). These results support the notion that H3-K27M-mutant cells may have already disrupted the MLL1 pathways, and thus are less sensitive to additional inhibition.

Gene Ontology (GO) analysis of genes associated with bivalent marks in SU-DIPG6 revealed enrichment for genes involved in neuron fate commitment and specification (Fig. S11E). Furthermore, RNA-seq revealed that genes associated with bivalent regions in H3-K27M mutant cultures are expressed at lower levels compared to WT cells (Fig. 5H). Thus, expression of mutant H3 not only re-patterns H3K27me3 and H3K4me3 marks, but also alters the highly balanced combinatorial code to generate an epigenetic state that facilitates gene silencing and may support an undifferentiated phenotype.

## Discussion

We applied high-resolution single-molecule imaging to explore chromatin incorporation of H3-K27M-mutant nucleosomes and their effect on the combinatorial epigenome of glioma cells. The single-molecule method is highly quantitative, enabling direct counting of mutant nucleosomes. It revealed differences in protein and incorporation levels between H3.1-K27M and H3.3-K27M nucleosomes, which may stem from the specific expression patterns of the two variants; H3.3-K27M chromatin incorporation occurs throughout the cell cycle, which may account for its higher total expression and fraction in chromatin (Tagami et al., 2004). Nevertheless, both variants show similar dramatic and global ramifications on the epigenetic landscape of these tumors, supporting epigenetic reprograming being one of the hallmarks of gliomagenesis (Phillips et al., 2020), and indicating a low threshold of expression for H3-K27M may suffice to drive chromatin alterations. Recent studies reported inhibition of PRC2 by peptides that mimic H3-K27M in their structure, and are not part of the chromatin (Jain et al., 2019; Piunti et al., 2019). Therefore, it remains possible that H3.3- and H3.1-mutant histones inhibit PRC2 also outside of the chromatin context, perhaps explaining the highly similar ramifications to H3K27me3 and H3K27ac for both mutant variants.

The combinatorial epigenetic analysis of mutant and WT nucleosomes point toward multiple nodes of epigenetic deregulation orchestrated by the H3-K27M oncohistone, with the activity of two critical epigenetic pathways disrupted, Polycomb and Trithorax. The single-molecule data supports increased binding between PRC2 and mutant nucleosomes in glioma cells, compatible with numerous biochemical assays (Bender et al., 2013b; Lewis et al., 2013a). The consequences of this increased binding, however, are dependent on the mutant histone variant, and specifically its chromatin environment. While canonical H3.1-K27M nucleosomes are indeed enriched for H3K27me3 on the WT histone, there is only mild enrichment for H3.3-K27M. Our results support the notion that this mild enrichment is a consequence of H3.3-K27M incorporation to active genomic regions by HIRA, presumably leading to the acquisition of active modifications that are known to inhibit EZH2 activity (Schmitges et al., 2011). In line with this notion, Stafford et al. reported that six hours following induced expression of H3-K27M in HEK293 cells, there is a transient enrichment of EZH2 on mutant nucleosomes, which is lost at later time points and replaced with H3K36me3 (Stafford et al., 2018). Overall, our analysis suggest that the combinatorial modification patterns of H3-K27M nucleosomes depend on both intrinsic features, such as binding affinity to chromatin modulators, and the functional features of the genomic region in which they reside.

We show that K27M-mutant nucleosomes directly interact with histone H3 lysine 4 methyltransferase, MLL1. Importantly, H3-K27M genomic distribution is not restricted to promoter regions (only 20% of peaks localize around TSS), thus its interaction with MLL1 leads to redistribution and modulation of H3K4me3 activity. Along with the acquisition of H3K4me3 in distal genomic regions, we provide evidence that the H3-K27M-MLL1 interaction may also contribute to upregulation of H3-K27M-specific transcriptional programs. The preferential incorporation of H3.3-K27M nucleosomes to actively transcribed genes may further strengthen the establishment of altered transcriptional programs. Furthermore, a key output of H3-K27M-mediated chromatin alterations to MLL1 and PRC2 is the increase in the fraction of bivalent chromatin, associated with low differentiation and high cellular plasticity, out of the regions marked with H3K27me3. Indeed, single-cell RNA sequencing in DIPG tumors showed most cells within the tumor resemble poorly differentiated oligodendrocyte precursor cells, with only a minority of malignant cells showing differentiated phenotypes (Filbin et al., 2018). While the altered bivalent state may at least partially stem from retention of H3K27me3 on CpG islands (Jain et al., 2020b), it may contribute to the dependency of these cells on residual EZH2 activity (Mohammad et al., 2017; Piunti et al., 2017). Interestingly, while we observe gain of bivalency in DIPG which is associated with gene repression, Larson et al. showed a different set of genes that lose bivalency and become activated, comprising an H3-K27M oncogenic signature (Larson et al., 2019). These results further support disruption of gene bivalency as a key epigenetic feature of glioma.

Our study highlights the utility of single-molecule tools to visualize the dynamic interplay between oncohistones and epigenetic pathways, thus revealing functional mechanisms by which tumorigenesis occurs. These advance tools will also be of significant value for the development and testing of new epigenetic-based therapeutic approaches.

## Acknowledgments

We thank H.Keren-Shaul, M.Pearl, O.Zion, R.Blecher (G-INCPM, WIS) for their help with HTS, H. Barr and A. Plotnikov for help with conducting the AlphaLISA assay, D.Deitch for graphical assistance, L. Segev for computational framework for single-molecule image analysis and I. Ulitsky for providing the pAG-MNase enzyme. We are thankful to N.Jabado for generously providing the H3-K27M and KO DIPG cultures (SU-DIPG13) (Krug et al., 2019). We are grateful for the important comments made by Y. Aylon, I. Tirosh and I. Ulitsky while reading the manuscript. Illustrations include content by BioRender.com.

## Author contributions

N.F., D.A and E.S designed the study and wrote the manuscript; N.F., D.A, O.B, N.M. performed experiments; B.D. performed bioinformatics analysis; V.F. adapted surface preparation protocols for single-molecule imaging; L.K. and S.B. generated and provided mouse NSC ChIP-seq data; M.M. provided tumor derived DIPG cells. D.J; contributed to the development of single-nucleosomes imaging technology described in this study.

## Competing interests

Authors declare no competing interests.

### Data and materials availability

All data is available in the main text or the supplementary materials. Sequencing data is deposited in NCBI’s Gene Expression Omnibus (GEO) and available through GEO series accession number GSE163184, GSE108364 and GSE171802.

### Supplementary Materials

Materials and Methods

Figures S1-S11

Table S1: Summary table of Cut&Run libraries

Table S2: Summary table of ATAC-seq libraries

Table S3: Summary table of MARS-seq libraries

