## Supplemental information for "H3-K27M-Mutant Nucleosomes Interact with MLL1 to Shape the Glioma Epigenetic Landscape"

### Supplementary information

Materials and Methods
Figures S1-S11
Table S1: Summary table of Cut&Run libraries
Table S2: Summary table of ATAC-seq libraries
Table S3: Summary table of MARS-seq libraries

### Materials and Methods

Cell culture:

All cell lines were maintained at 37˚C with 5% CO2. HEK293 cells were cultured in DMEM supplemented with 10% FBS and 1% P/S. DIPG derived cells, SU-DIPG-6 (H3.3-K27M), SU-DIPG-13 (H3.3-K27M), SU-DIPG-17 (H3.3-K27M), SU-DIPG-36 (H3.1-K27M), SU-DIPG-38 (H3.1-K27M) and SU-DIPG-48 (H3 WT) were generated in the lab of Dr. Michelle Monje, Stanford University and grown as previously described (Grasso et al., 2015). K27M-KO SU-DIPG13 and corresponding control clone were generated in the lab of Prof. Nada Jabado, McGill University, as previously described (Krug et al., 2019). Briefly, cells were cultured in Tumor Stem Media (TSM) consisting of a 1:1 mixture of DMEM/F12 (Invitrogen) and Neurobasal (-A) (Invitrogen), with addition of B27(-A) (Invitrogen), human-bFGF (20 ng/ml) (Shenandoah Biotechnology), human-EGF (20 ng/ml) (Shenandoah Biotechnology), human PDGF-AA (20 ng/ml) (Shenandoah Biotechnology), human PDGF-BB (20 ng/ml) (Shenandoah Biotechnology) and heparin (10 ng/ml) (Stemcell Technologies). Cells were passaged every week.

Generation of doxycycline inducible cell lines:

pInducer20 H3.3 WT and H3.3-K27M were a gift from M. Suva. H3.1 WT plasmid was generated by RF cloning and site directed mutagenesis was used to generate the H3.1-K27M plasmid. Cloned plasmids were validated by Sanger sequencing. Lentiviral (3$^{rd}$ generation) packaging was performed by jetPEI-mediated transfection of HEK293T with appropriate plasmids. Virus-containing supernatants were collected 48h following transfection, filtered, supplemented with 8µg/ml Polybrene, and added to HEK293 target culture. Infected cells were selected with 0.5mg/ml G418 and induced with 1 µg/ml doxycycline.

siRNA transfection:

For siRNA-mediated knockdown, the indicated SMARTpools (Dharmacon) were used with Lipofectamin2000 transfection reagent according to the manufacturer`s instructions. Final siRNA concentration was 20nM in all cases. The medium containing oligonucleotides and reagents was replaced after 6h.

FLAG-H3 plasmid transfection:

pcDNA4/TO-Flag-H3.3 plasmid was obtained from Addgene (#47980) and site-directed mutagenesis was used to introduce K27M mutation. jetPEI-mediated transfection was conducted according to manufacturer instructions and cells were collected after 72h.

Cell viability analysis

Cells were plated at a density of 1250 cells per well in 96-well plates in at least triplicate and subjected to drug treatment for 5-8 days. When treated for 8 days one pulse were given after 96h. Cell viability was measured by CelltiterGlo assay (G7571, Promega) according to the manufacturer's instructions. Luminescence was measured by Cytation 5 plate reader and viability was compared to DMSO treated control cells.

Nucleosome preparation for single-molecule imaging:

2-2.5 million cells were collected and washed with PBS supplemented with protease inhibitors cocktail (1:100, Sigma P8340), and HDAC inhibitors (20mM Sodium butyrate, Sigma 303410 and 0.1mM Vorinostat V-8477), and centrifuged at 3000 rpm for 3 minutes. Cell pellet was resuspended with 1ml of 0.05% IGEPAL (Sigma I8896) diluted in PBS (supplemented with inhibitors) and centrifuged once again at 3000 rpm for 3 minutes. Pellet was then resuspended in Lysis buffer (100mM Tris-HCl pH 7.5, 300mM NaCl, 2% Triton® X-100, 0.2% sodium deoxycholate, 10mM CaCl2) supplemented with inhibitors and Micrococcal Nuclease (ThermoFisher Scientific, 88216). The suspension was incubated at 37°C for 10 minutes and then inactivated by addition of EGTA at a final concentration of 20mM. Next, the lysate was centrifuged for 10 min at max speed and the supernatant was transferred to a new tube. 10% of the lysate was used to extract DNA by AMPure SPRI beads (Beckman Coulter, A63881) and chromatin digestion was verified by resolving on 2% Agarose gel. Nucleosomes were then concentrated on an Amicon ultra-4 (Millipore, UFC810024) for 20 minutes followed by 1 minute centrifugation at 16,000g. Inhibitors were supplemented following concentration.
For nucleosomes labeling reaction mix contained: NEBuffer™ 2 (NEB B7202), protease and HDAC inhibitors (as detailed above), 0.25 mM MnCl2, 33uM fluorescently labeled dATP (Jena Bioscience, NU-1611-Cy3/Cy5), 33uM biotinylated dUTP (Jena Bioscience , NU-803-BIOX), 1.5ul of Klenow Fragment (3'→5' exo-, NEB, M0212S) and 1.5ul of T4 Polynucleotide Kinase (NEB, M0201L). Sample was incubated at 37°C for 1 hour and then inactivated by addition of EDTA at a final concentration of 20mM. Nucleosomes were then purified on Performa Spin Columns (EdgeBio, 13266) followed by the addition of inhibitors. To verify nucleosome labeling, 10ul of the nucleosomes and 10ul of ANPure extracted DNA, were analyzed on 6% TBE gel (ThermoFisher Scientific, EC62655BOX) and imaged with Typhoon imager (Amersham Biosciences).

Surface preparation for single-molecule imaging

PEG-biotin microscope slides were prepared based on the protocol described by Chandradoss et al. (Chandradoss et al., 2014). Ibidi glass coverslips (25 mm x 75 mm, IBIDI, IBD-10812) were cleaned with (1) MilliQ H2O (3X washes, 5 min sonication, 3X washes), (2) 2% Alconox (Sigma 242985) (20 min sonication followed by 5X washes with MilliQ H2O), (3) 100%

Acetone (20 min sonication followed by 3X washes with MilliQ H2O). To ensure surface functionality, slides were incubated in 1M KOH solution for 30 min while sonicated (Sigma 484016), followed by 3X washes with MilliQ H2O. Slides were sonicated for 10 min in 100% HPLC ethanol (J.T baker 8462-25) prior to applying amino-silanization chemistry. Slides were incubated for 24 min in a mixture of 3% 3-Aminopropyltriethoxysilane (ACROS Organics, 430941000) and 5% acetic acid in HPLC EtOH), with 1 min sonication in the middle. Slides were then washed with HPLC EtOH (3X) and MilliQ H2O (3X) and dried with nitrogen. The first step of passivation was performed by applying mPEG:biotin-PEG solution (20mg Biotin-PEG (Laysan, Biotin-PEG-SVA-5000), 180mg mPEG (Laysan, MPEG-SVA-5000) dissolved in 1560ul 0.1M Sodium Bicarbonate (Sigma, S6297) on one surface followed by the assembly of another surface on top. Each pair of assembled surfaces were incubated overnight in a dark humid environment. At the next day, surfaces were washed with MilliQ H2O and dried with N2 followed by the second passivation step. MS (PEG) 4 (ThermoFisher Scientific, TS-22341) was diluted in 0.1M of sodium bicarbonate to a final concentration of 11.7 mg/ml and applied on one surface, followed by the assembly of another surface on top. Each pair of assembled surfaces were incubated overnight in dark humid environment, washed with MilliQ H2O and dried with nitrogen. After Nitrogen flush, surfaces are stored in -20°C.

Single-molecule imaging

PEG-biotin coated coverslips were assembled into Ibidi flowcell (Sticky Slide VI hydrophobic, IBIDI, IBD-80608). Streptavidin (SIGMA, S4762) was added to a final concentration of 0.2mg/ml followed by an incubation of 10 minutes. TetraSpeck beads (ThermoFisher Scientific, T7279) diluted in PBS were added and incubated on surface for at least 30 minutes. Labeled nucleosomes were incubated for 10 minutes in imaging buffer (10mM MES pH 6.5 (Boston Bioproducts Inc, NC9904354), 60mM KCL, 0.32mM EDTA, 3mM MgCl2, 10% glycerol, 0.1mg/ml BSA (Sigma, A7906), 0.02% Igepal (Sigma, I8896)) to allow immobilization via biotin-streptavidin interactions and washed with imaging buffer. Antibodies were diluted in imaging buffer to a final concentration of 50-100ng/ml and incubated for 30 minutes. All positions (80-100 fields-of-view per experiment) were then imaged by a total internal reflection (TIRF) microscope by Nikon (Ti2 LU-N4 TIRF) every 10 minutes (10-15 cycles).

Image analysis

Image analysis was performed with Cell Profiler image analysis tools (http://www.cellprofiler.org/). Briefly, image analysis is done in three steps: (1) Time-lapse images of antibody binding events and TetraSpeck beads are aligned, stacked and summed to one image. Antibody spots and TetraSpeck beads spots were distinguished based on the size of the spot. (2) Stacked images are aligned to the initial images of the nucleosomes based on TetraSpeck beads location spots, and only binding events that align with nucleosomes are filtered and saved for further analysis. To evaluate random co-localization (negative control), each stacked image is aligned to a 90° flipped image of the initial nucleosomes. For H3-K27M images, all imaged nucleosomes were divided into either K27M-mutant or WT nucleosomes according to the summed antibody signal. (3) Filtered modified nucleosomes for each single mark are aligned to identify combinatorial marks on single nucleosomes. Average percentage of modified nucleosomes over all fields of view (FOV) imaged ± SE is shown. Since we use the TetraSpeck beads and the microscope stage is highly accurate, alignment of images results in shifts of up to 30 pixels. Nucleosomes are initially distributed on the surface in low density to minimize overlap between spots.

Chromatin bound assay

Chromatin-Bound fraction isolation was conducted as previously described (Shema et al., 2011) and whole cell lysate and the chromatin fraction were resolved by SDS-PAGE.

Cut&Run pulldown assay followed by high-throughput sequencing

Cut&Run assay was done as described in (Meers et al., 2019; Skene and Henikoff, 2017) with slight modifications as follows: Cells were harvested and counted, with 200,000 cells taken per reaction. Permeabilized cells, bound to Concanavalin A-coated beads, were mixed with individual primary antibody (see antibody list) and incubated overnight at 4°C while rotating. Secondary antibody, anti-rabbit HRP, was used as a negative control. pAG-MNase enzyme (generated in the Department of Life Sciences Core Facilities, WIS, using Addgene plasmid 86973) was added to each sample followed by incubation step of 1 hour at 4°C. Targeted digestion was done by 15 minutes incubation on ice block (0°C) under low salt conditions. DNA purification was done using Nucleospin gel and PCR clean-up kit (Machery-Nagel, 740609).

Libraries were prepared from 1-20ng of DNA as previously described(Blecher-Gonen et al., 2013). Libraries were quantified by Qubit (ThermoFisher Scientific) and TapeStation (Agilent). Sequencing was done on a Next-Seq 500 instrument (Illumina) using a V2 150 cycles mid output kit, allocating 10M reads per sample (paired end sequencing).

Reads were preprocessed with cutadapt (Martin, 2011) to remove adapters and low-quality bases (parameters: --times 2 -q 30 -m 20), and reads quality was evaluated using FastQC. Reads were mapped to human genome (hg19, UCSC) using Bowtie version 2.3.5.1 (Langmead and Salzberg, 2012) (--local --very-sensitive-local --no-unal --no-mixed --no-discordant -I 10 -X 700). Nucleosome fragments at the length >120bp were selected from the remaining unique reads using picard-tools, and broad peaks were called using MACS2 against the corresponding HRP samples as background control (parameters: -f BAMPE --SPMR --nomodel --extsize 100 --keep-dup auto -q 0.05). MLL1 peaks were called using narrow peaks. Bigwig files were constructed from BAM alignments using deepTools2 suite (Ramírez et al., 2016) with bamCoverage, using RPKM-normalization in 10bp bins, and excluding ENCODE hg19 blacklist regions. Heatmaps and profiles were constructed in 'scale-regions' mode around peak summits, with missingDataAsZero paramter, and were clustered using the K-means method in 'plotHeatmap'. Spearman correlation coefficients were computed using 'plotCorrelation'. Peak intersections were computed with bedtools. RPKM-normalized coverage files were visualized on the genome using IGV (2.8.6). Analysis of genomic features was done using ChIPseeker, with promoters defined as +/- 3kb from TSS (Yu et al., 2015). Annotation of CpG islands were downloaded for UCSC. Association of peaks with specific genes and functional enrichment of groups of peaks was done using GREAT (McLean et al., 2010). Reads coverage around TSS was visualized using ngs.plot (Shen et al., 2014). Enhancers were defined as genomic regions consisting H3K27ac peaks and do not overlap within +/-2Kb of known TSS.

ATAC-seq library preparation and sequencing

Sample preparation was conducted as previously described by Buenrostro et al (Buenrostro et al., 2015), with modifications described by Lara-Astiaso and colleges (Lara-Astiaso et al., 2014). Briefly, 50,000 cells from each culture were used, and each cell type was analyzed in two replicates. Nuclei were incubated with 2ul of Nextera Tn5 enzyme (TDE1, Illumina) for 1h at 37°C. Enzyme inactivation was done by addition of 5ul Clean-up buffer (900mM NaCl, 30mM EDTA), 2ul of 5% SDS and 2ul of Proteinase K (NEB) and incubation for 30min at 40°C. Tagmentated DNA was isolated using 2x SPRI beads cleanup.

For library amplification, two sequential 9-cycle and 5-cycle PCR were performed in order to enrich small tagmentated DNA fragments. Libraries were prepared using KAPA HiFi HotStart ready mix. After the first PCR, the libraries were selected for small fragments using SPRI cleanup (0.65x). Then a second PCR was performed with the same conditions in order to obtain the final library. DNA concentration was measured with a Qubit fluorometer (Life Technologies) and library sizes were determined using TapeStation (Agilent Technologies). Libraries were sequenced on the NovaSeq6000 sequencing platform using SP, 100cycles kit (paired end sequencing), with an average of 140 million reads obtained for each sample.

Paired-end raw reads were trimmed using cutadapt (Martin, 2011) (with the parameters -q 25 -m 30), and mapped to the human genome hg19 (hg19, UCSC) using bowtie2 (Langmead and Salzberg, 2012) (with the parameters --local). Following alignment, mitochondrial genes were removed from the analysis, and duplicated reads were removed using picard-tools.

Nucleosome-free fragments at the length <120bp were selected from the remaining unique reads, and broad peaks were called using MACS2 (with the parameters --bw 120 -B -f BAMPE --SPMR –B --shift -50 --extsize 100 -keep-dup all -q 0.05). DeepTools2 (Ramírez et al., 2016) was used to generate profiles, with missingDataAsZero parameter.

Isolation of total RNA and Bulk MARS-Seq library preparation and sequencing.

RNA was isolated using the NucleoSpin kit (Macherey Nagel). A bulk adaptation of the MARS-Seq protocol (Jaitin et al., 2014; Keren-Shaul et al., 2019) was used to generate RNA-Seq libraries for expression profiling. Briefly, 30 ng of input RNA from each sample was barcoded during reverse transcription and pooled. Following Agencourct Ampure XP beads cleanup (Beckman Coulter), the pooled samples underwent second strand synthesis and were linearly amplified by T7 in vitro transcription. The resulting RNA was fragmented and converted into a sequencing-ready library by tagging the samples with Illumina sequences during ligation, RT, and PCR. Libraries were quantified by Qubit and TapeStation as well as by qPCR for GAPDH housekeeping gene as previously described. Sequencing was done on a Nextseq 75 cycles high output kit (Illumina). An average of 15 million reads were obtained for each sample.

MARS-seq analysis was preformed using the UTAP transcriptome analysis pipeline (Kohen et al., 2019). Reads were trimmed to remove adapters and low quality bases using cutadapt (Martin, 2011) and mapped to the human genome (hg19, UCSC) using STAR v2.4.2a (Dobin et al., 2012) (parameters: –alignEndsType EndToEnd, –outFilterMismatchNoverLmax 0.05, –twopassMode Basic, –alignSoftClipAtReferenceEnds No). The pipeline quantifies the 3' of annotated genes (The 3' region contains 1,000 bases upstream of the 3' end and 100 bases downstream). Counting was done using HTSeq-count (Anders et al., 2014) in union mode. Genes having a minimum 5 UMI-corrected reads in at least one sample, were considered. Normalization of the counts and differential expression analysis was done using DESeq2 (Love et al., 2014) (parameters: betaPrior=True, cooksCutoff=FALSE, independentFiltering=FALSE). Differentially expressed genes were defined as genes that had a significant adjusted p-value (with Benjamini and Hochberg procedure) <= 0.05, log2FoldChange >= 1 and baseMean >= 5. H3.3-K27M up regulated gene signature were defined as genes that were significantly upregulated in both SU-DIPG6 and SU-DIPG13 compared to SU-DIPG48. Expression levels are shown as log2 of the average normalized count from the 3 replicates. Only genes that had an average of >=5 normalized counts in all samples examined, are presented in the box-plots. Expression heat-map was generated using Morpheus (https://software.broadinstitute.org/morpheus). Genes listed in table S6 by Filbin and colleges(Filbin et al., 2018) was used as the H3-K27M-specific gene signature shown in figures 5E-F.

Mouse Hindbrain NSC Chromatin Immunoprecipitation and sequencing (ChIP-seq)

Mice strains generation, data acquisition and sequencing analysis were described in by Larson et al. (Larson et al., 2019). Data processing for H3K27me3 ChIPseq was also as described in Larson et al. Drosophila melanogaster spike in reads were cleaned from H3K4me3 ChIP and inputs using bbmap 38.86 bbduk and bbsplit with a hybrid mm9/dm6 genome. H3K4me3, FLAG and input fastqs were processed with the nf-core/chipseq v1.2.1 pipeline with BWA v0.7.17 -r1188 for mapping to mm9, BEDTools v2.29.2 bedGraphToBigWig for bigwig file generation and MACS2 v2.2.7.1 for peak calling using the –narrow_peak and –macs_fdr 0.01 flags. DeepTools2 suite was used to generate heatmaps and profiles. Peak intersections were computed with bedtools. Analysis of genomic features was done using ChIPseeker. H3K4me3 antibody is Cell Signaling 9571 (lot 8) and FLAG antibody is Sigma F1804 (lot SLBJ4607V).

Data availability

Cut&Run, ATAC and MARS-seq data is deposited in NCBI's Gene Expression Omnibus (GEO) and available through GEO series accession number GSE158447.

ChIP-seq data from mouse NSC is available at GSE108364 (H3K27me3) and GSE171802 (H3K4me3 and FLAG).

Protein-protein binding assay

Biotinylated nucleosomes (1µg) (Epicypher; 16-0006, 16-0317, 16-1323) were immobilized on 10 µl BSA-blocked Dynabeads™ M-280 Streptavidin beads (Invitrogen, ThermoFisher Scientifics) in 0.01% BSA/PBS supplemented with protease inhibitors (1:100, Sigma, P8340) for 1 hour on ice. The beads were then incubated with 0.2ug Recombinant EZH2 protein complex (ActiveMotif, 31337) for 2 hours at 4°C. Complexes were washed with 0.01% BSA/PBS, 0.1% BSA/PBS, and 0.1% BSA, 0.05% Tween/PBS (3X). After washes the protein-bound beads were re-suspended in protein sample buffer and analyzed by western blotting with anti-H3 and anti EZH2 antibodies.

For AlphaLISA protein-protein interaction assay 10ul of biotinylated nucleosomes (10nM) were incubated with 10ul of Recombinant KMT2A (MLL1) complex (50nM, ActiveMotif, 31423) or EZH2 complex (2.5nM, ActiveMotif, 31337) for 2 hours in binding buffer (0.5% BSA, 0.01% NP4 in PBS). 10ul of Glutathione or Anti-6xHis AlphaLISA Acceptor Beads (80ug/ml, PerkinElmer, AL109C, AL178C) were added to each well in a 96 well ½ area Alpha plate (PerkinElmer, 6002350) and incubated protected from light for 2 hours, followed by addition of 10ul AlphaScreen Streptavidin Donor beads (PerkinElmer, 6760002S) and additional 2 hours incubation. Signal was measured by PHERAstar FS microplate reader (615 nm, AlphaLISA filter).

Histone Methyltransferase Assay

H3K4me3 methyltransferase assay was conducted using EpiQuik™ Histone Methyltransferase Activity/Inhibition Assay Kit (H3-K4) (Epigentek). Biotinylated substrate was replaced with biotinylated recombinant nucleosomes (Epicypher).

Statistics

Unless noted otherwise, p-values were determined using one or two-tailed, two-sample t-tests.

Antibodies:

| Antibody | Vendor | Identifier |
|---|---|---|
| Histone H3 (K27M Mutant Specific) (D3B5T) Rabbit mAb | Cell signaling | CST-74829S |
| Histone H3 (K27M Mutant Specific) (D3B5T) Rabbit mAb (Alexa Fluor 488 Conjugate) | Cell signaling | CST-74829S custom conjugation |
| Peroxidase-AffiniPure Goat Anti-Rabbit IgG | Jacksoon Immuno Research Laboratory | 111-035-144 |
| Tri-Methyl-Histone H3 (Lys27) (C36B11) Rabbit mAb (Alexa Fluor 647 Conjugate) | Cell signaling | CST-12158 |
| Acetyl-Histone H3 (Lys9) (C5B11) Rabbit mAb (Alexa Fluor 647 Conjugate) | Cell signaling | CST-4484 |
| Tri-Methyl-Histone H3 (Lys4) (C42D8) Rabbit mAb (Alexa Fluor 555 Conjugate) | Cell signaling | CST-11960 |
| Acetyl-Histone H3 (Lys27) (D5E4) XP® Rabbit mAb | Cell signaling | CST-8173P |
| Tri-Methyl-Histone H3 (Lys27) (C36B11) Rabbit mAb (Alexa Fluor 488 Conjugate) | Cell signaling | CST-5499S |
| Ezh2 (D2C9) XP(R) Rabbit mAb | Cell signaling | CST-5246S |
| Acetyl-Histone H3 (Lys27)(D5E4) XP(R) Rabbit mAb (Alexa Fluor(R) 647 Conjugate) | Cell signaling | CST-39030S |
| Tri-Methyl-Histone H3(K27) (C36B11) Rabbit mAb | Cell signaling | CST-9733S |
| Tri-Methyl-Histone H3(K4) (C42D8) Rabbit mAb | Cell signaling | CST-9751S |
| MLL1 (D6G8N) Rabbit mAb (Carboxy-terminal Antigen) | Cell signaling | CST-14197S |
| Histone H3 (D1H2) XP® Rabbit mAb | Cell signaling | CST-4499 |
| Anti-Histone H3.3 antibody [EPR17899] | Abcam | AB-ab208690 |
| MONOCLONAL ANTI-FLAG(R) M2-CY3 antibody | SIGMA | A9594 |

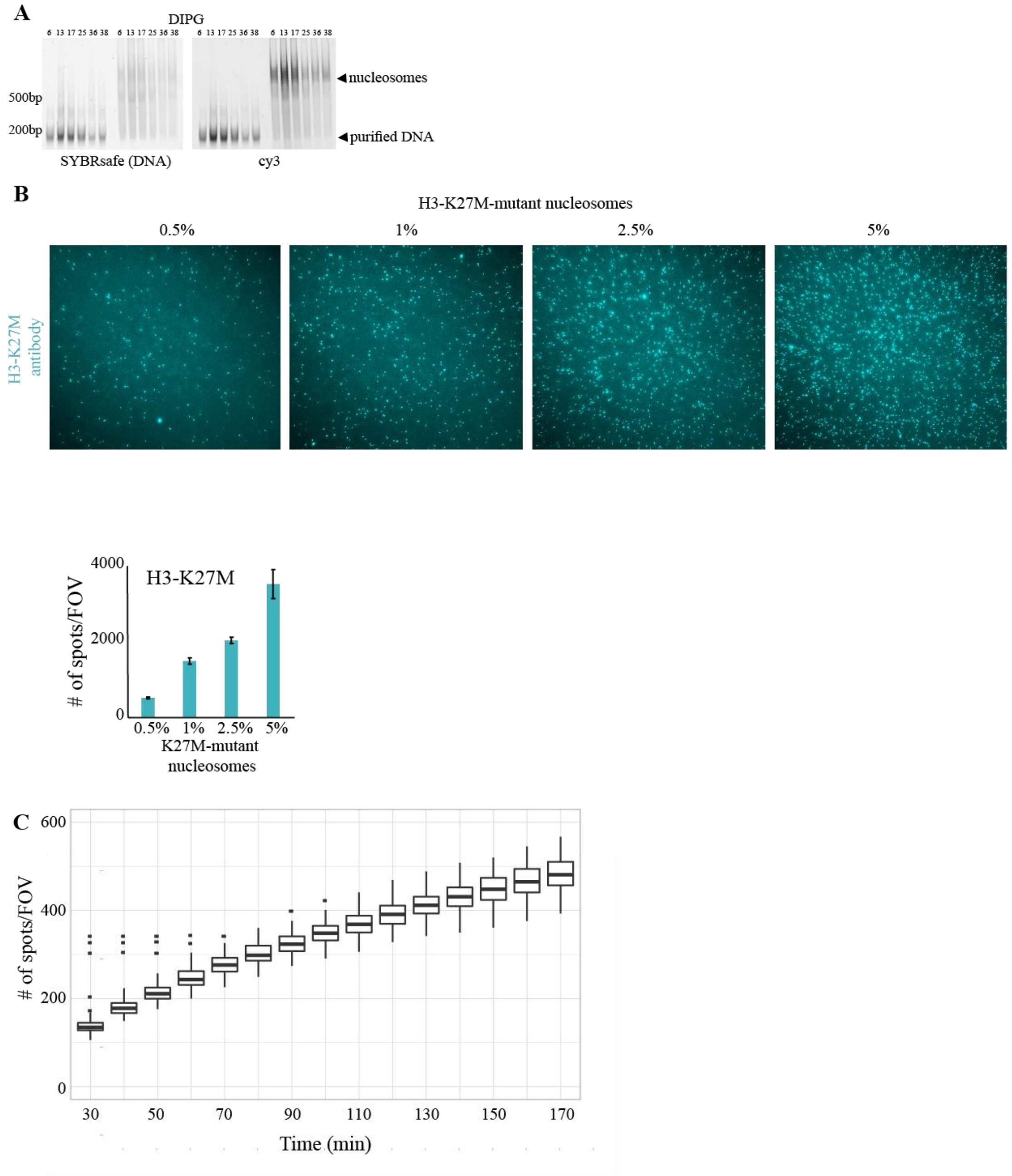


**Figure S1: (A)** Preparation of nucleosome samples for single-molecule imaging: mono-nucleosomes were extracted from the indicated DIPG cell lines and labeled with cy3-dATP (see methods). Both purified DNA and intact nucleosomes were resolved on a 6% TBE gel and visualized by Typhoon laser scanner (cy3 and cy2 channels are shown). **(B)** Recombinant H3-K27M-mutant nucleosomes were mixed with WT nucleosomes at the indicated ratios and quantified by single-molecule imaging using H3-K27M-AF488 antibody. Top: representative fields-of-view (FOVs) from each sample. Bottom: quantification of all FOVs imaged. **(C)** Representative quantification of the cumulative signal obtained from imaging nucleosomes extracted for HEK293 cells expressing H3.3-K27M. Imaging the same FOV over time allows detection of maximal antibody events.

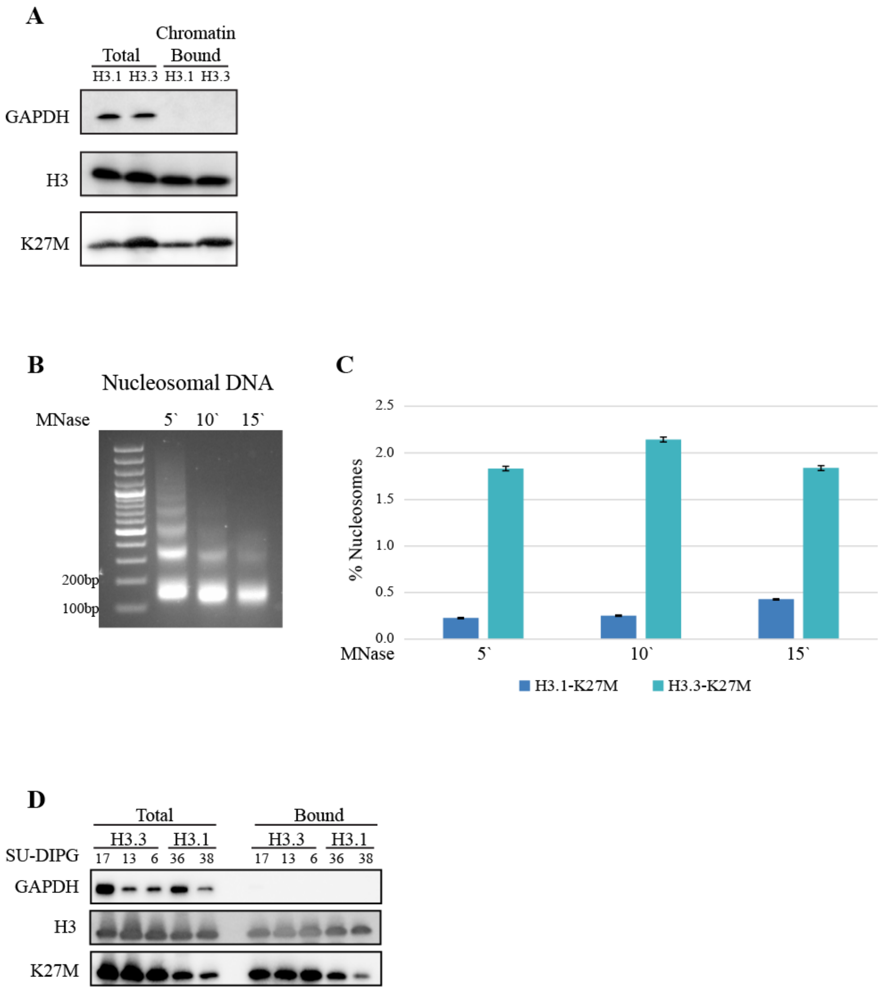


**Figure S2: (A)** Whole cell lysate (total) and chromatin bound fraction from HEK293 cells expressing H3.1- or H3.3-K27M in a doxycycline dependent manner were analyzed by western blot using the indicated antibodies. **(B)** Chromatin from HEK293 cells was treated with MNase for the indicated times (minutes), resulting in different degrees of digestion. Shorter incubation results in larger portion of the chromatin remaining at higher ordered structures, and release of mono-nucleosomes from more accessible genomic regions. **(C)** Single-molecule imaging and quantification of H3-K27M-mutant nucleosomes in H3.1- or H3.3-K27M HEK293 cells, treated with MNase as described in B. Higher levels of H3.3-K27M compared to H3.1-K27M are observed in the various MNase digestion conditions. **(D)** SU-DIPG cells were analyzed as in A. Higher levels of H3.3-K27M nucleosomes compared to H3.1-K27M nucleosomes are observed in the DIPG lines.

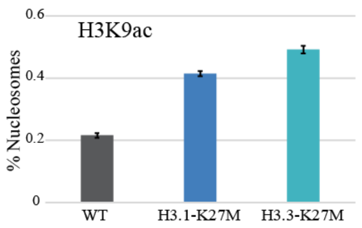


**Figure S3:** Single-molecule imaging and quantification of H3K9ac-modified nucleosomes in HEK293 cells expressing H3-K27M. Higher levels of acetylation is observed in cells harboring the H3-K27M-mutant histone compared to cells expressing WT H3.

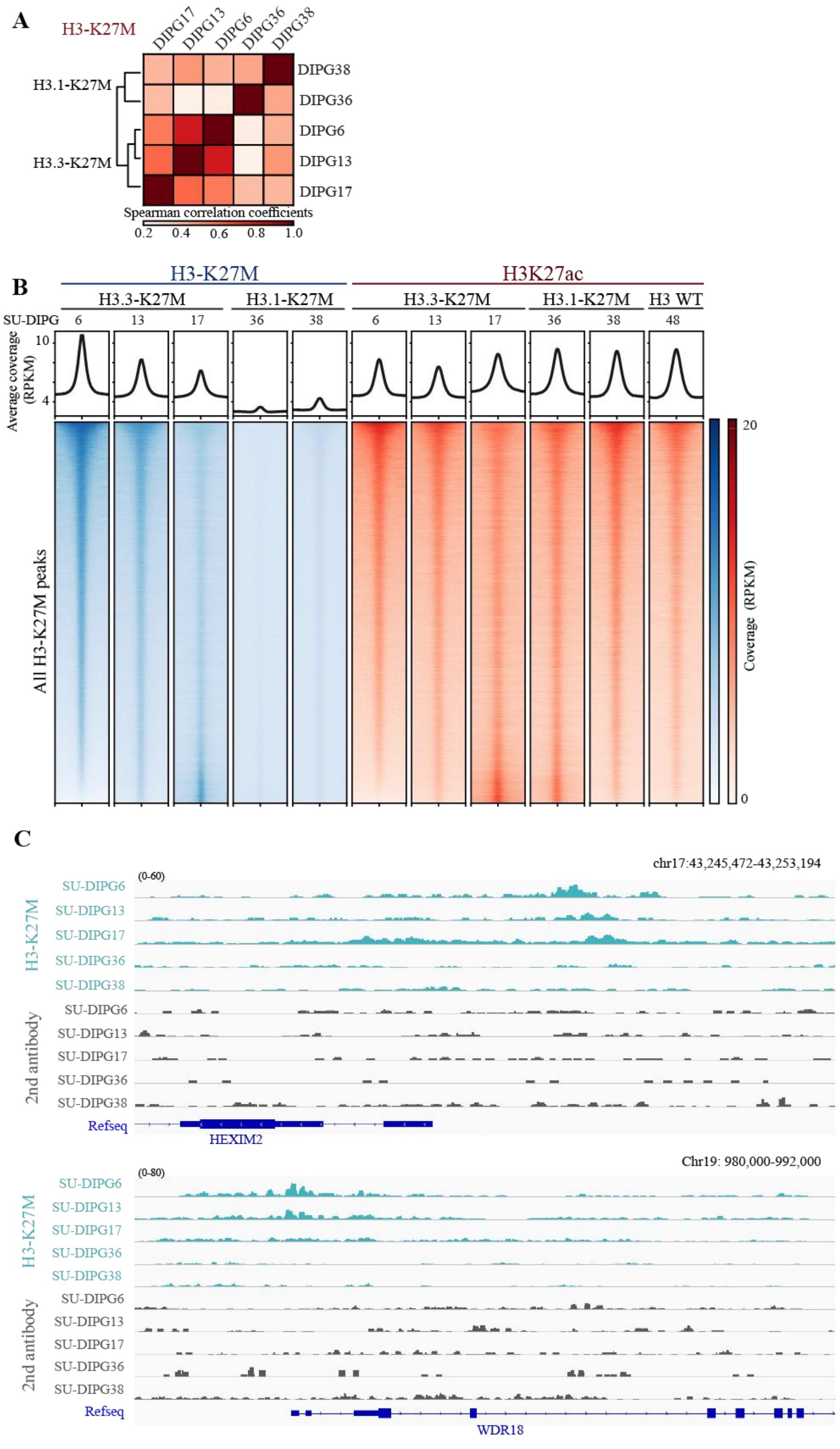

A
H3-K27M
DIPG17
DIPG13
DIPG6
DIPG36
DIPG38
H3.1-K27M
H3.3-K27M
Spearman correlation coefficients
0.2 0.4 0.6 0.8 1.0
B
H3-K27M
H3K27ac
H3.3-K27M
H3.1-K27M
H3 WT
SU-DIPG
6 13 17 36 38 6 13 17 36 38 48
Average coverage (RPKM)
10
4
All H3-K27M peaks
Coverage (RPKM)
20
0
C
chr17:43,245,472-43,253,194
(0-60)
H3-K27M
SU-DIPG6
SU-DIPG13
SU-DIPG17
SU-DIPG36
SU-DIPG38
2nd antibody
Refseq
HEXIM2
Chr19: 980,000-992,000
(0-80)
WDR18

D

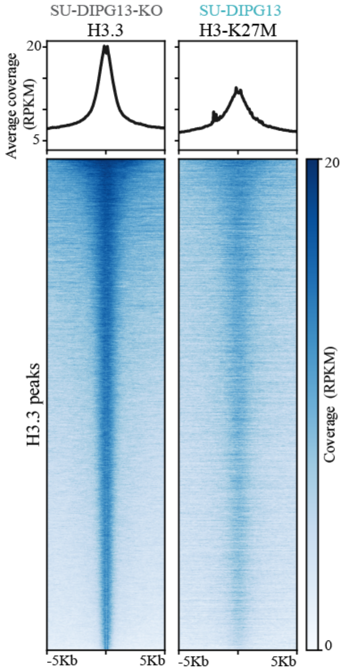


**Figure S4: (A)** Overall similarity between DIPG-lines based on H3-K27M reads' coverage (Spearman correlation coefficients). Higher correlation is observed between cells with H3-K27M mutation on histone variant H3.3. **(B)** Cut&Run using H3-K27M and H3K27ac antibodies in the indicated patient-derived DIPG cultures, followed by high-throughput sequencing. H3-K27M peaks detected in all cultures analyzed (n=121,298) are shown in the heatmap, sorted in descending order from top to bottom according to SU-DIPG6 (normalized counts (RPKM) are shown ±5kb around the peak summit). The signal obtained for H3-K27M and H3K27ac at these regions is plotted accordingly for the indicated samples. Average coverage is plotted on top. Similar incorporation patterns are observed for H3.3-K27M lines, as opposed to SU-DIPG36 and SU-DIPG38 harboring H3.1-K27M which do not form peaks. **(C)** Representative IGV genome browser tracks for H3-K27M peaks and the corresponding signal obtained for secondary antibody only (negative control). **(D)** Heatmap of H3.3 peaks identified in SU-DIPG13 knocked out for H3.3-K27M (thus, the signal corresponds only to WT H3.3), plotted along with the corresponding signal of H3-K27M in the matched isogenic line (expressing H3.3-K27M). Signal is sorted according to H3.3 intensity.

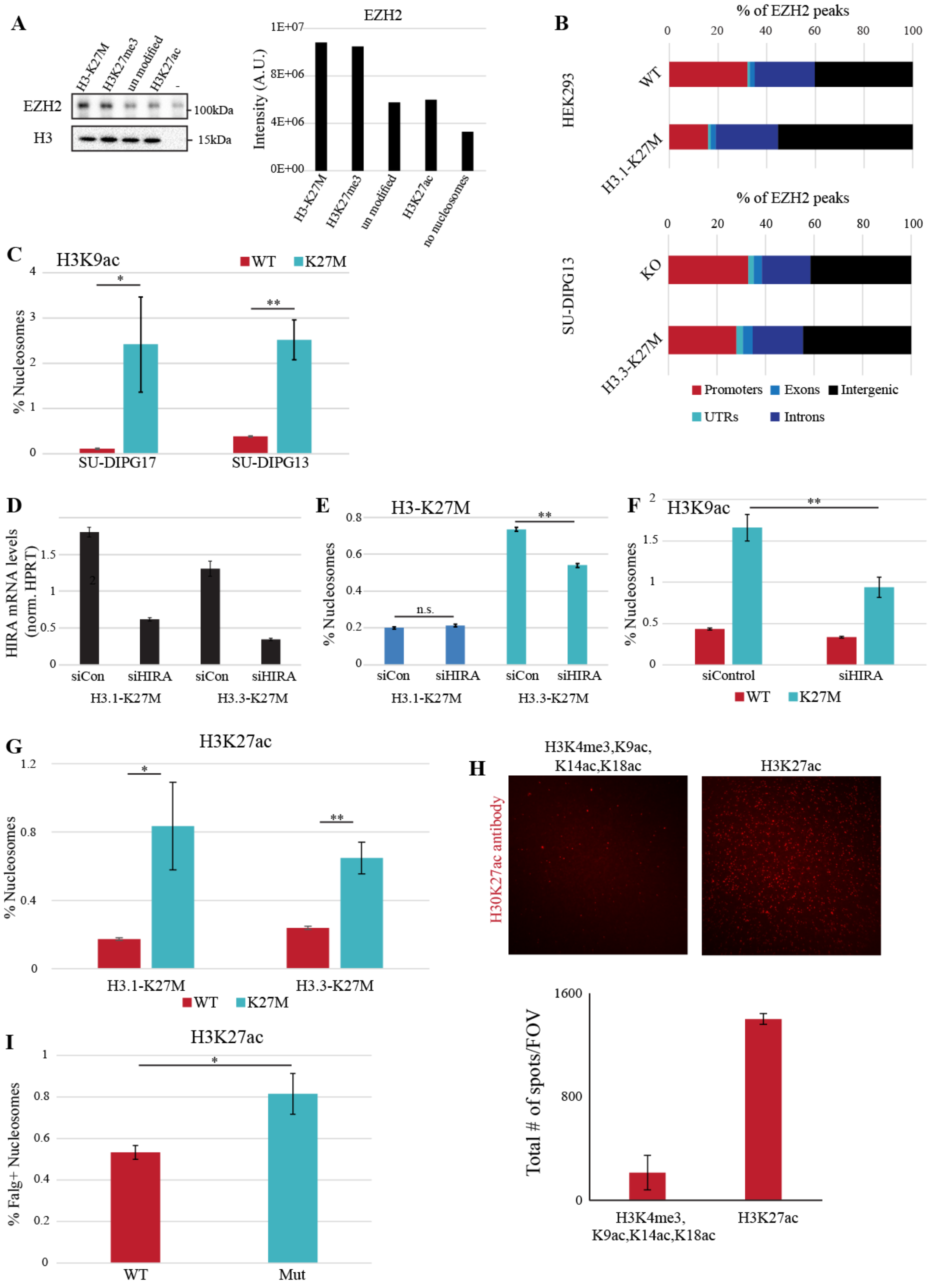


**Figure S5: (A)** Preferential binding of PRC2 to H3-K27M nucleosomes. Binding assay between recombinant EZH2 protein complex and biotinylated recombinant nucleosomes (H3-K27M, H3K27me3, H3K27ac or unmodified WT nucleosomes). Following incubation with streptavidin-coated beads, complexes were washed, resolved on SDS-PAGE and visualized using the indicated antibodies. Quantification of EZH2 signal intensity is shown on the right. **(B)** Proportion of EZH2 peaks that correspond to the indicated genomic features in HEK293 cells expressing either WT or K27M-mutant H3.1 (up) and SU-DIPG13 knocked out for H3.3-K27M and the appropriate control cells (bottom). Expression of H3-K27M results in higher proportion of EZH2 peaks associated with distal genomic regions. **(C)** Single-molecule analysis of H3K9ac on H3-K27M-mutant nucleosomes in the indicated DIPG lines. Percentage of H3K9ac nucleosomes out of WT and H3-K27M positive nucleosomes is shown. *p-val<0.05, **p-val<0.001. **(D-F)** HEK293 cells were transfected with the indicated siRNAs and the expression of either H3.1-K27M or H3.3-K27M histones was induced by doxycycline. Cells were collected after 72 hours, knockdown efficiency was evaluated by qRT-PCR **(D)**, and H3-K27M nucleosomes were quantified by single-molecule imaging. **(E).** HIRA depletion leads to reduced chromatin incorporation of H3.3-K27M, with no effect on H3.1-K27M. **(F)** Nucleosomes from the indicated samples were analyzed as in B. Depletion of HIRA results in reduction of H3-K27M-H3K9ac nucleosomes. **(G)** Single-molecule analysis of H3K27ac on H3-K27M-mutant nucleosomes in HEK293 cells expressing H3.1- or H3.3-K27M. Percentage of wild type (WT) or H3-K27M-mutant nucleosomes acetylated on lysine 27. Enrichment of H3K27ac on mutant nucleosomes (i.e. heterotypic nucleosomes) is observed for both variants. **(H)** The indicated recombinant nucleosomes were analyzed by single-molecule imaging using H3K27ac antibody. Each modified nucleosome was mixed with un-modified recombinant nucleosomes to obtain a mixture of 5% modified nucleosomes. Upper panel: representative FOV from each sample. Lower pannel: quantification of all FOVs imaged. **(I)** HEK293 cells were transfected with plasmids encoding FLAG-H3.3 or FLAG-H3.3-K27M and nucleosomes were imaged using anti-FLAG and H3K27ac antibodies. Percentage of H3K27ac marked nucleosomes out of FLAG positive nucleosomes is shown.

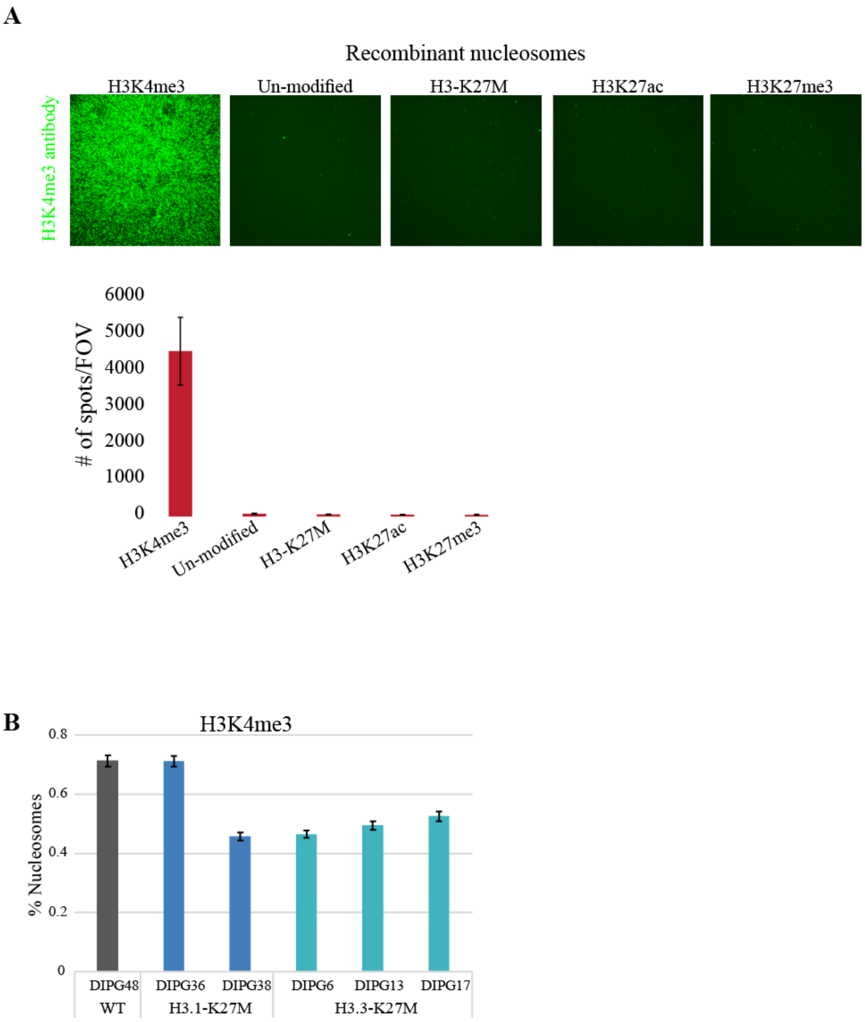


**Figure S6: (A)** The indicated recombinant nucleosomes were analyzed by single-molecule imaging using H3K4me3 antibody. Representative FOVs are shown in the upper panel and quantification of all FOVs imaged is shown at the bottom panel. **(B)** Single-molecule quantification of H3K4me3 nucleosomes in the indicated DIPG cells lines.

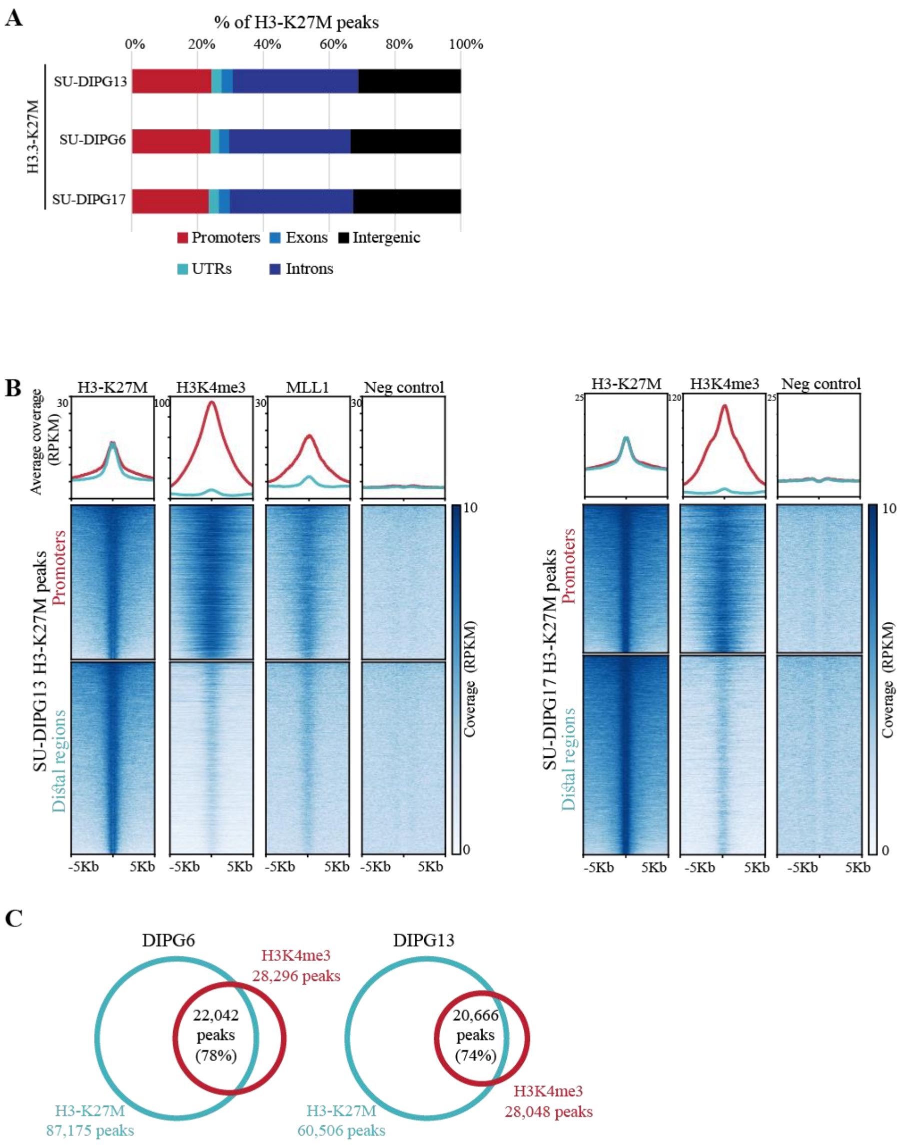


**Figure S7: (A)** Proportion of H3-K27M peaks that correspond to the indicated genomic features in the indicated DIPG cultures. **(B)** H3-K27M peaks identified in SU-DIPG13 (Left) and SU-DIPG17 (right) cells are associated with H3K4me3 and MLL1 in both promoters and distal genomic regions. Heatmap shows H3-K27M peaks annotated to promoters (±3Kb from TSS) or distal genomic regions using ChIPseeker algorithm sorted in descending order from top to bottom (center represents the peak summit, left and right borders represent -5kb and +5kb, respectively). Signal for H3K4me3, MLL1 (for SU-DIPG13 only) and negative control antibody (anti-rabbit 2$^{nd}$ antibody) in these genomic regions is plotted accordingly. Average coverage for each group is plotted on top. **(C)** Peak intersection analysis for H3-K27M and H3K4me3 peaks in the indicated DIPG cells. Intersecting peaks have at least 1bp overlap. Percentage is calculated out of H3K4me3 peaks.

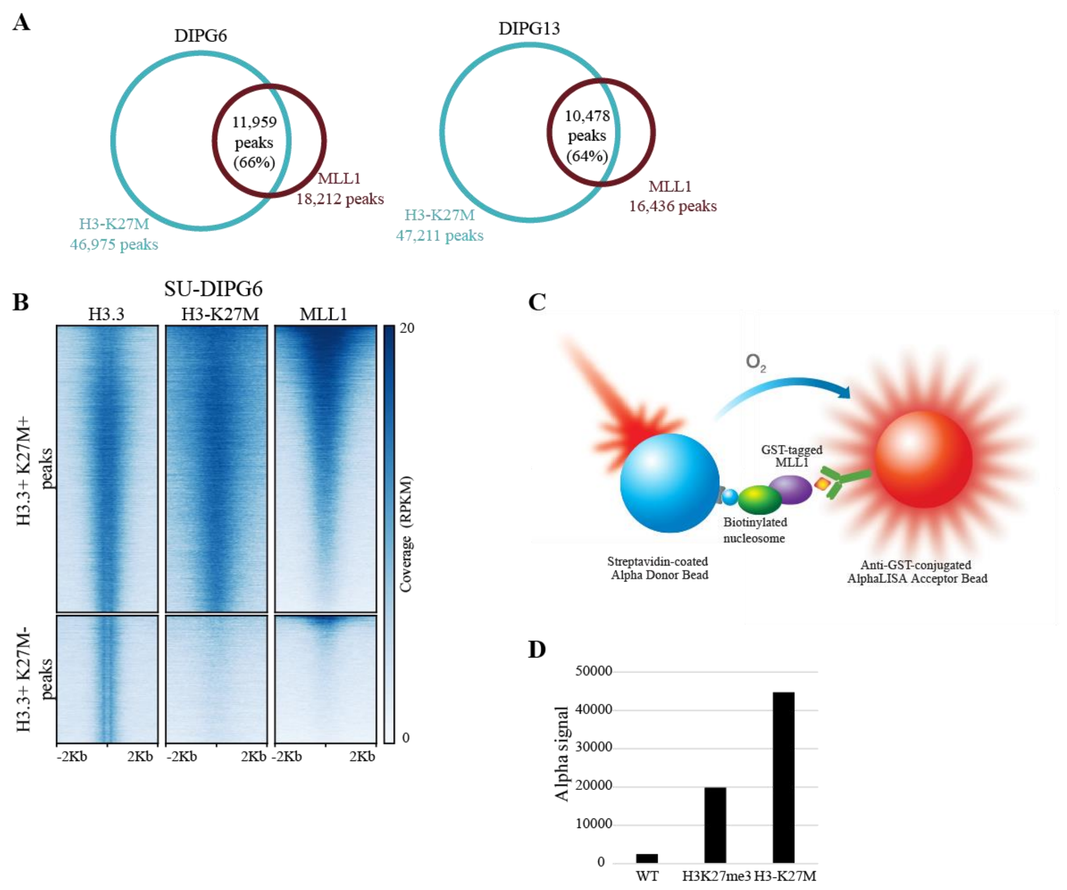


**Figure S8: (A)** Peak intersection analysis for H3-K27M and MLL1 peaks in the indicated DIPG cells. Intersecting peaks have at least 1bp overlap. Percentage is calculated out of MLL1 peaks. **(B)** H3.3 peaks identified in Cut&Run analysis of H3.3-K27M DIPG culture (SU-DIPG6) were divided according to the overlap with H3-K27M peaks, and the corresponding signal of MLL1 in these regions is shown. For each group, regions are sorted according to MLL1 signal. Results indicate that MLL1 is associated with genomic regions that are positive to both H3.3 and H3-K27M. (**C)** Illustration of an Alpha protein-protein interaction assay: one protein is captured on the donor beads, and the other protein is captured on the acceptor beads. When the two proteins interact, the donor bead is brought into proximity of the acceptor bead, and excitation of the donor bead will result in signal generation dependent on the presence of an interaction (adapted from the assay user guide, PerkinElmer). **(D)** AlphaLISA protein-protein binding assay indicates preferential binding of EZH2 to H3-K27M-mutant and H3K27me3 nucleosomes over WT nucleosomes. 2.5nM of GST-tagged MLL1 was incubated with 10nM of biotinylated recombinant nucleosomes. Alpha acceptor and donor beads were added, and the Alpha signal corresponding to protein-protein binding was measured.

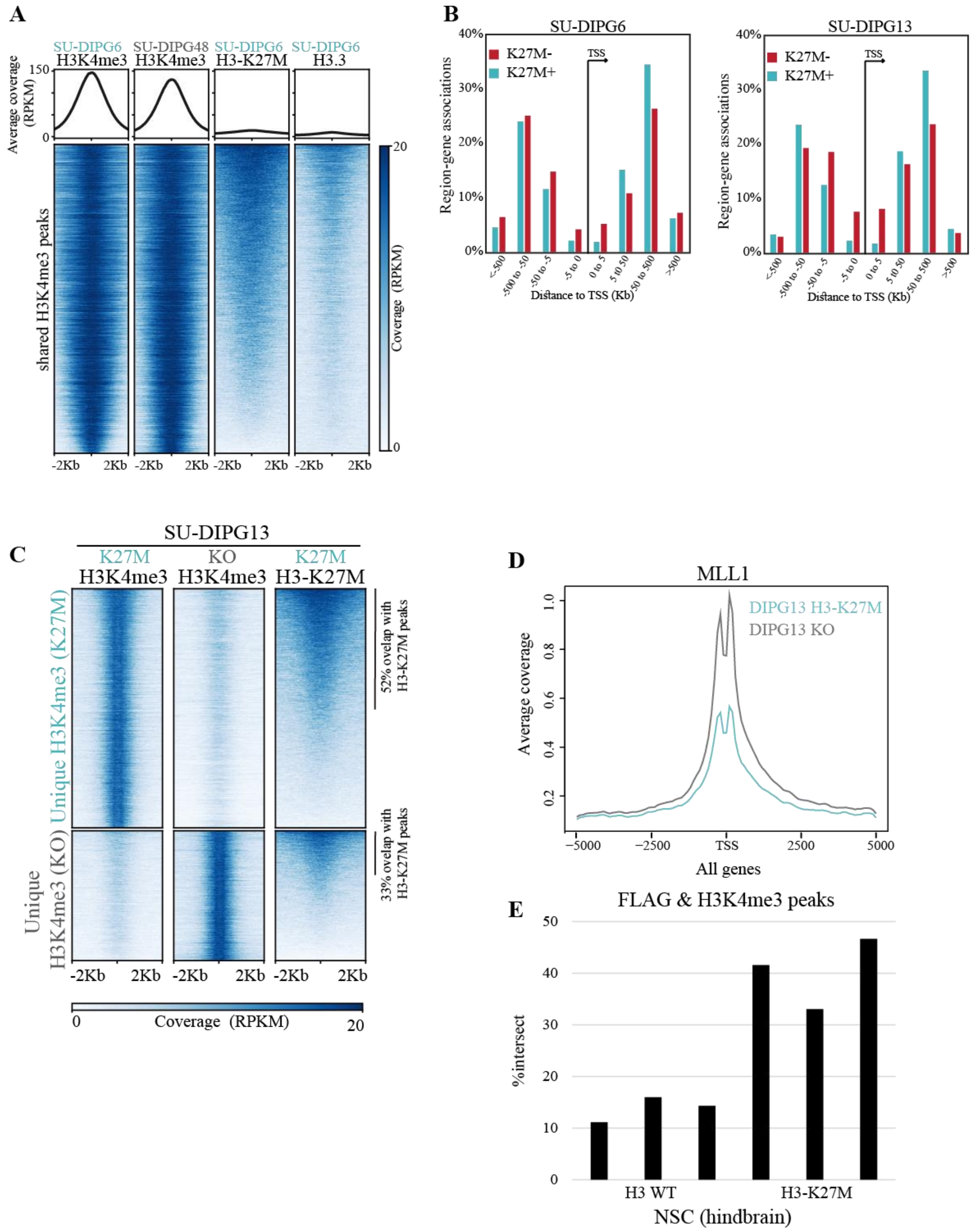


**Figure S9: (A)** Heatmap of shared H3K4me3 peaks, as defined in figure 3B (10,499 peaks). **(B)** H3K4me3 unique peaks were divided according to their overlap with H3-K27M peaks. GREAT algorithm was used to associate the peaks with genes depending on the distance from the nearest TSS. Percentage region-gene associations for each distance is shown. **(C)** Similar heatmap as shown in figure 3E, including both H3K4me3 peaks that are unique to H3.3-K27M or KO cells (center represents the peak summit, left and right borders represent -5kb and +5kb, respectively). H3-K27M signal in these genomic regions is plotted accordingly and the overlap with identified H3-K27M is noted for each group. H3-K27M cells show a larger number of unique H3K4me3 peaks and 52% of these peaks are associated with H3-K27M peaks, as compared to only 33% of KO-unique H3K4me3 peaks which localize to regions marked with H3-K27M in the mutant cells. **(D)** MLL1 coverage (read count per million mapped reads) over TSS in H3.3-K27M-mutant and K27M-KO isogenic cells (SU-DIPG13). **(E)** ChIP-seq using FLAG and H3K4me3 antibodies in hindbrain NSC expressing FLAG tagged WT or K27M-mutant H3.3. Percentage of H3K4me3 peaks that overlap with FLAG peaks (minimal overlap of 25%) from three mice from each genotype.

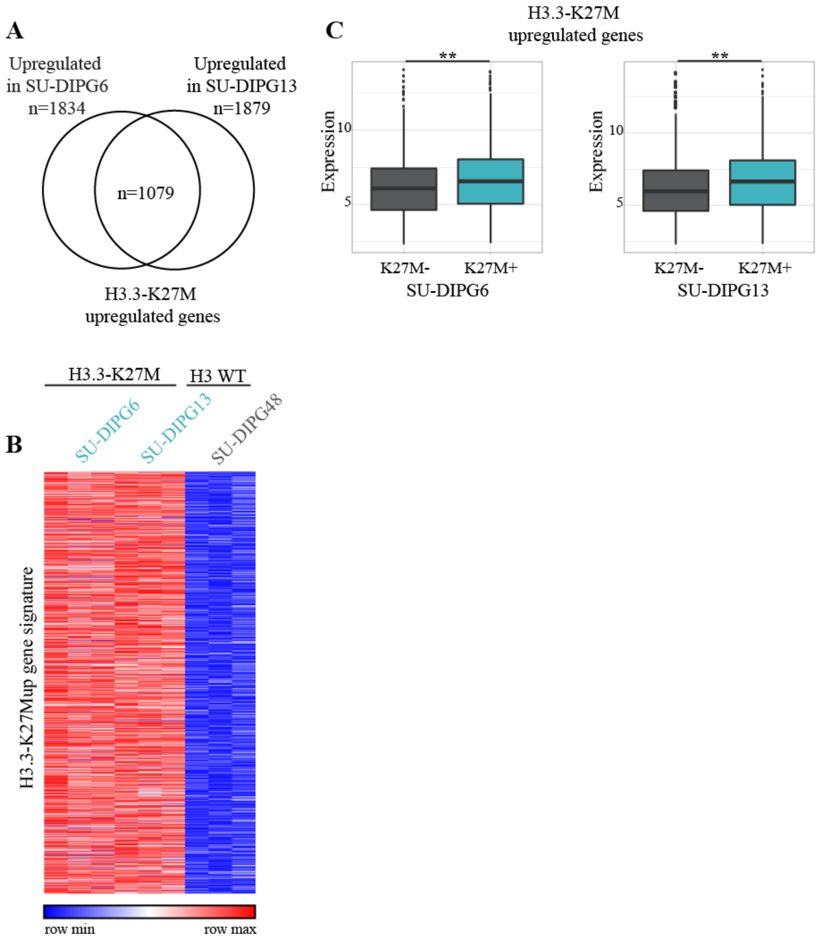


**Figure S10: (A)** H3.3-K27Mup gene signature consist of genes significantly upregulated in both SU-DIPG6 and SU-DIPG13 when compared to WT H3 cells, SU-DIPG48 (upper panel). **(B)** Heatmap depicting the expression levels of H3.3-K27Mup gene signature in the indicated DIPG cultures. Standardized rld values for three replicates of each culture are shown. **(C)** H3.3-K27M upregulated genes were divided according to the presence of H3-K27M peak ±2Kb form their TSS (as in figure 5D-E), and the average expression levels for each group of genes is shown in SU-DIPG6 cells (left) and SU-DIPG13 (right).

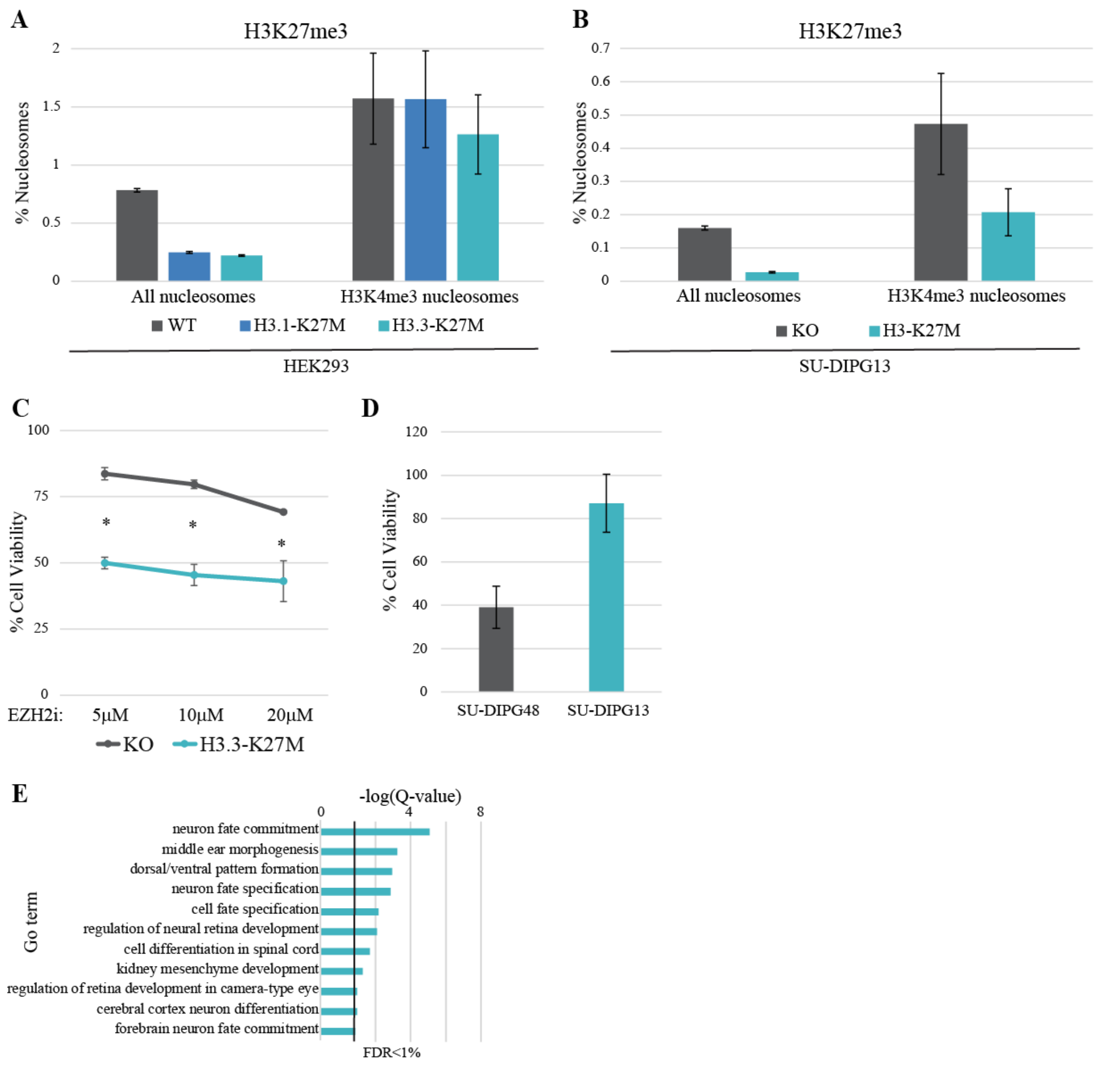


**Figure S11: (A-B)** Alternative presentation of the single-molecule analysis of bivalent nucleosomes shown in fig. 5E-F. Single nucleosomes marked with H3K27me3 or H3K4me3 were identified. Percentage of H3K27me3+ nucleosomes out of all nucleosomes, or only out of H3K4me3+ nucleosomes, is shown. While H3-K27M leads to strong reduction in the number of H3K27me3 nucleosomes, H3K4me3+ nucleosomes retain the H3K27me3 mark. **(C)** SU-DIPG13 knocked out for H3-K27M (KO) and their corresponding H3-K27M expressing cells were treated with EZH2 inhibitor (EPZ6438) at the indicated concentrations. Cell viability was measured by CellTiterGlo assay relative to DMSO treated cells (mean ± SE for three technical replicates, *p-val<0.05). **(D)** SU-DIPG13 (H3.3-K27M) and SU-DIPG48 (WT-H3) were treated with 30µM MLL1 inhibitor (MM-102). Cell viability was measured by CellTiterGlo assay relative to DMSO treated cells. Mean ± SE are shown for two technical repeats). **(E)** Functional enrichment analysis of genes associated with bivalent peaks (+/- 1kb from TSS) identified in SU-DIPG6 cells.

**Table S1.**

Summary table of Cut&Run libraries

| Cell line | Antibody | No. of uniquely mapped paired-end reads | % Uniquely and concordantly aligned reads | No. of broad peaks (against HRP control) | No. of narrow peaks (against HRP control) |
|---|---|---|---|---|---|
| SU-DIPG13 - R1 | **HRP** | 20,560,709 | 72.45 | | |
| | **K27M** | 33,647,503 | 82.03 | 78,115 | |
| | **H3K27ac** | 28,922,557 | 83.28 | 61,205 | |
| | **H3K27me3** | 21,112,951 | 75.21 | 22,921 | |
| | **H3K4me3** | 31,522,380 | 85.25 | 40,584 | |
| SU-DIPG13 – R2 | **HRP** | 10,022,093 | 67.54 | | |
| | **H3K27ac** | 17,032,339 | 83.51 | 65,609 | |
| | **K27M** | 14,241,430 | 81.64 | 60,506 | |
| | **H3K27me3** | 13,730,774 | 75.89 | 16,722 | |
| | **H3K4me3** | 16,315,702 | 86.52 | 28,048 | |
| SU-DIPG17-R1 | **HRP** | 7,657,863 | 71.30 | | |
| | **H3K27ac** | 25,758,695 | 84.77 | 80,437 | |
| | **K27M** | 31,989,874 | 82.46 | 37,043 | |
| | **H3K27me3** | 12,011,679 | 72.59 | 16,917 | |
| | **H3K4me3** | 8,522,480 | 85.01 | 23,417 | |
| SU-DIPG36-R1 | **HRP** | 3,848,292 | 68.95 | | |
| | **H3K27ac** | 21,730,163 | 86.80 | 45,177 | |
| | **K27M** | 16,596,066 | 76.56 | 29 | |
| | **H3K27me3** | 8,893,757 | 75.09 | 6,809 | |
| | **H3K4me3** | 8,559,652 | 86.36 | 24,348 | |
| SU-DIPG38-R1 | **HRP** | 11,676,341 | 69.82 | | |
| | **H3K27ac** | 8,752,948 | 85.13 | 58,614 | |
| | **K27M** | 14,995,272 | 77.43 | 1,737 | |
| | **H3K27me3** | 9,201,420 | 70.88 | 13,018 | |
| | **H3K4me3** | 10,120,163 | 85.46 | 22,597 | |

| | | | | | |
|---|---|---|---|---|---|
| | **HRP** | 11,678,133 | 69.64 | | |
| | **H3K27ac** | 9,417,273 | 84.20 | 63,258 | |
| | **K27M** | 10,489,332 | 82.47 | 87,175 | |
| | **H3K27me3** | 11,793,389 | 73.09 | 20,730 | |
| **SU-DIPG6-R1** | **H3K4me3** | 14,587,881 | 82.00 | 28,296 | |
| | **HRP** | 12,068,023 | 70.04 | | |
| | **H3K27ac** | 13,430,897 | 82.36 | 45,299 | |
| | **H3K27me3** | 11,262,733 | 74.27 | 90,187 | |
| **SU-DIPG48-R1** | **H3K4me3** | 14,183,765 | 82.00 | 22,889 | |
| | **HRP** | 9,152,411 | 68.95 | | |
| | **K27M** | 7,752,347 | 81.56 | 47,211 | |
| **SU-DIPG13-R3** | **MLL1** | 8,066,930 | 74.29 | | 16,436 |
| | **HRP** | 13,180,732 | 69.49 | | |
| | **H3K27me3** | 16,057,285 | 75.04 | 86,337 | |
| **SU-DIPG48-R2** | **MLL1** | 10,301,424 | 72.51 | | 16,544 |
| | **HRP** | 4,317,903 | 69.48 | | |
| | **K27M** | 2,781,211 | 83.38 | 46,975 | |
| **SU-DIPG6-R2** | **MLL1** | 3,247,903 | 75.05 | | 18,212 |
| | **K27M** | 18,425,336 | 96.24 | 64,509 | |
| | **MLL1** | 26,890,124 | 92.77 | | 35,207 |
| | **H3K4me3** | 33,287,560 | 94.92 | 27,540 | |
| | **HRP** | 28,473,038 | 87.99 | | |
| **SU-DIPG6-R3** | **H3.3** | 37,656,602 | 91.09 | 41,157 | |
| | **K27M** | 8,187,636 | 83.25 | 73,078 | |
| | **H3K4me3** | 10,907,782 | 85.81 | 37,435 | |
| | **MLL1** | 8,077,502 | 75.48 | | 20,813 |
| | **HRP** | 12,154,980 | 71.91 | | |
| **SU-DIPG13-K27M** | **H3K27me3** | 11,831,965 | 77.30 | 16,631 | |
| **SU-DIPG13-KO** | **H3K4me3** | 11,927,867 | 87.70 | 32,107 | |

| | | | | | |
|---|---|---|---|---|---|
| | **HRP** | 4,986,262 | 68.62 | | |
| | **MLL1** | 8,305,829 | 76.57 | | 23,011 |
| | **H3.3** | 10,649,998 | 77.88 | 34,031 | |
| | **H3K27me3** | 9,360,655 | 80.86 | 51,375 | |
| **HEK293-H3.1WT** | **EZH2** | 10,427,930 | 78.81 | 31,077 | |
| | **HRP** | 7,330,406 | 70.98 | | |
| **HEK293-H3.1K27M** | **EZH2** | 10,666,046 | 78.40 | 35,016 | |
| | **HRP** | 9,061,569 | 72.19 | | |

**Table S2.**

Summary table of ATAC-seq libraries

| Cell line | Replicate | No. of reads (total) | No. of uniquely mapped paired-end reads | % Uniquely and concordantly aligned reads | Number of nucleosome-free reads (properly paired) | No. of broad Peaks |
|---|---|---|---|---|---|---|
| **SU-DIPG13** | 1 | 144,644,480 | 61,261,196 | 42.80% | 24,633,642 | 94,118 |
| | 2 | 166,263,642 | 78,276,591 | 47.44% | 54,258,760 | 94,990 |
| **SU-DIPG48** | 1 | 135,711,822 | 77,681,957 | 57.71% | 89,683,094 | 117,696 |
| | 2 | 120,178,240 | 67,207,762 | 56.40% | 71,812,726 | 121,584 |
| **SU-DIPG6** | 1 | 114,496,573 | 62,712,282 | 55.05% | 45,712,206 | 144,596 |
| | 2 | 172,044,560 | 90,063,067 | 52.72% | 61,324,054 | 141,087 |

**Table S3.**

Summary table of MARS-seq libraries

| Cell line | Replicate | No. of reads (total) | No. of Uniquely mapped reads | % of Uniquely mapped reads | No. of reads uniquely mapped to genes | No. of reads after UMI correction |
|---|---|---|---|---|---|---|
| SU-DIPG6 | 1 | 24,082,492 | 14,653,912 | 70.59 | 6,893,019 | 3,422,635 |
| | 2 | 26,645,718 | 15,453,290 | 70.95 | 6,000,074 | 2,952,929 |
| | 3 | 15,761,902 | 9,638,735 | 72.28 | 7,884,123 | 3,826,948 |
| SU-DIPG13 | 1 | 21,521,770 | 13,123,001 | 70.73 | 6,869,606 | 3,530,475 |
| | 2 | 13,370,601 | 8,059,645 | 71.35 | 9,563,237 | 4,392,994 |
| | 3 | 13,565,171 | 8,204,046 | 71.27 | 6,314,864 | 2,921,148 |
| SU-DIPG36 | 1 | 20,154,339 | 12,179,371 | 71.06 | 8,639,999 | 4,415,886 |
| | 2 | 14,075,735 | 8,432,517 | 70.68 | 6,689,082 | 3,510,389 |
| | 3 | 16,891,071 | 9,801,923 | 70.08 | 4,456,323 | 2,296,799 |
| SU-DIPG38 | 1 | 23,714,452 | 13,895,732 | 69.95 | 6,724,465 | 3,466,970 |
| | 2 | 16,543,610 | 9,493,850 | 70.7 | 5,320,202 | 2,726,140 |
| | 3 | 10,817,873 | 6,493,997 | 71.96 | 4,894,630 | 2,403,209 |
| SU-DIPG48 | 1 | 17,275,879 | 10,536,142 | 72.85 | 5,775,805 | 2,991,745 |
| | 2 | 19,081,221 | 11,631,547 | 73.18 | 6,640,424 | 3,193,494 |
| | 3 | 16,501,100 | 10,008,787 | 71.96 | 8,785,916 | 4,194,507 |
